# A structural and regulatory framework for Atg9-containing vesicle formation and their Atg1-dependent remodelling during autophagy initiation

**DOI:** 10.1101/2025.11.06.685569

**Authors:** Carmen Oeo-Santos, Ella Knüpling, Colin Davis, Xin Meng, Sarah Maslen, Simone Kunzelmann, Rocco D’Antuono, Anna Olerinyova, Tania Auchynnikava, Mark Skehel, Anne Schreiber

**Author notes:** These authors contributed equally (Carmen Oeo-Santos, Ella Knüpling).

## Abstract

Autophagy is a complex intracellular degradation pathway that depends on the coordinated interplay between the core autophagy machinery and diverse membrane sources to drive the *de novo* formation of double-membrane vesicles, known as autophagosomes. Golgi-derived Atg9-containing vesicles are essential for this process, delivering membranes to the pre-autophagosomal structure (PAS). These vesicles contain the transmembrane proteins Atg9 and Atg27 and the peripheral membrane protein Atg23; however, the nature, function, and regulation of their interactions remain poorly understood.

Here, we systematically dissect the molecular interactions between Atg9, Atg23 and Atg27, and uncover their regulation in space and time. The bipartite binding mode by which Atg23 engages Atg9 provides a structural model for how Atg23 promotes vesicle budding. Furthermore, Atg1-dependent phosphorylation of Atg9 remodels its interactions with Atg23 and Atg27 at the PAS to support autophagy initiation. Together, these findings establish a molecular and regulatory framework for the earliest steps of autophagy.

## Introduction

Macroautophagy, hereafter referred to as autophagy, is a highly conserved intracellular degradation pathway that utilizes *de novo* double-membrane vesicle (autophagosome) formation to engulf cytoplasmic material and deliver it to the lysosome or vacuole for degradation and subsequent recycling. While bulk autophagy indiscriminately captures cytoplasmic material, selective autophagy pathways specifically engulf a range of targets, including damaged organelles, protein aggregates, and intracellular pathogens. Autophagy is therefore critical for maintaining cellular homeostasis, particularly under nutrient-limiting conditions, and for supporting cellular quality control and cell intrinsic defence mechanisms by preventing the accumulation of potentially cytotoxic material. Dysregulation of autophagy is implicated in cellular aging and has been linked to various human diseases, including cancer, neurodegeneration, and infectious disorders^1^. Consequently, modulating autophagy presents a promising therapeutic strategy for treating a wide spectrum of diseases and to potentially increase health span.

Autophagosome formation is a complex process requiring the coordinated interplay of the core autophagy machinery and various membrane and lipid sources^2–4^. Atg9-containing vesicles are thought to be the key membrane source initiating autophagosome biogenesis^5–7^. These vesicles bud from the Golgi apparatus and shuttle to the pre-autophagosomal structure (PAS), the site of autophagosome formation^7,8^, where they are tethered by one of the key regulators of autophagy, the Atg1 complex^9,10^. Potential fusion of these Atg9-containing vesicles, together with localized lipid synthesis^11^, Atg2-mediated lipid shuttling^12,13^ and phospholipid transfer between the cytoplasmic and inner membrane leaflets catalysed by Atg9’s lipid scramblase activity^14^, contribute to the emergence and expansion of a curved membrane sheet, known as the isolation membrane or phagophore, which upon closure gives rise to a double-membrane autophagosome.

Atg9 is a multi-span transmembrane protein that is essential for both bulk and selective autophagy^15^. Its transmembrane region has been shown to form a homotrimer^14,16–19^ with its long, largely disordered N- and C-terminal regions both facing the cytoplasm^20^. In *Saccharomyces cerevisiae (S. cerevisiae)*, these cytoplasmic regions mediate the interactions with the bulk and selective autophagy-specific Atg1 complex subunits Atg17 and Atg11, allowing the tethering of Atg9-containing vesicles at the PAS^9,10,21^. The serine/threonine protein kinase Atg1, which forms part of both the bulk and selective Atg1 complexes, has been shown to phosphorylate the cytoplasmic regions of Atg9^22–24^. This posttranslational regulation is required to promote autophagy^23,24^, although the underlying molecular mechanism remains largely elusive.

In addition to Atg9, *S. cerevisiae* Atg9-containing vesicles also contain the single-span transmembrane protein Atg27 and the peripheral membrane protein Atg23^25,26^. Atg23 and Atg27 both play a key role in bulk autophagy and are critical for most selective autophagy pathways^26,27^. Deletion of either Atg23 or Atg27 impairs Atg9-containing vesicle formation and, as a result, their recruitment to the PAS^6,17,27–29^. However, it remains unclear whether these two proteins only regulate Atg9-containing vesicle formation or also their tethering. While Atg9 has been suggested to interact with Atg23^26^ and Atg27^25^ via its cytoplasmic N- and C-terminal tails, no interaction could be detected between Atg23 and Atg27 in the absence of Atg9^28^. It remains to be investigated whether the interactions of Atg9 with Atg23 and Atg27 are direct, and how they are regulated at the molecular level.

To elucidate the earliest steps of autophagosome formation, we analysed the organization of Atg9-containing vesicles using *in vitro* reconstitution, structure prediction, mass spectrometry, and a combination of biochemical, biophysical, and cell biology approaches. By systematically elucidating the molecular basis of the protein–protein interactions between key vesicle components and their regulation, we gained insights into the formation and organization of Atg9-containing vesicles and their Atg1-dependent remodelling at the PAS, a key step in autophagy initiation.

## Results

### Atg23 interacts with both the cytoplasmic N- and C-terminal regions of Atg9

To investigate Atg9’s cytoplasmic interaction network with the core Atg9-containing vesicle components Atg23 and Atg27 (**Fig. 1a**) in a detergent-free environment that preserves protein-protein interactions, we generated an Atg9 construct lacking only the transmembrane region (TMR). Firstly, we tested the resultant construct, Atg9^ΔTMR^ (**Fig. 1b**), by assessing its interactions with the bulk and selective autophagy-specific Atg1 complex subunits Atg17 and Atg11, respectively. Both proteins have previously been shown to mediate the recruitment of Atg9 to the PAS^30^. In agreement with previous studies^9,31^, we could show that the Atg9^ΔTMR^ directly interacts with Atg11 (**Extended Data Fig. 1a**). In addition, Atg9^ΔTMR^ formed a stoichiometric complex with Atg17 (**Extended Data Fig. 1b**). Consistent with previous findings^10^, Atg9^ΔTMR^-Atg17 complex formation could be abolished by addition of the Atg29-Atg31 subcomplex (**Extended Data Fig. 1b**), further validating Atg9^ΔTMR^ as a suitable tool for studying Atg9’s cytoplasmic protein-protein interactions.

**Fig. 1:**
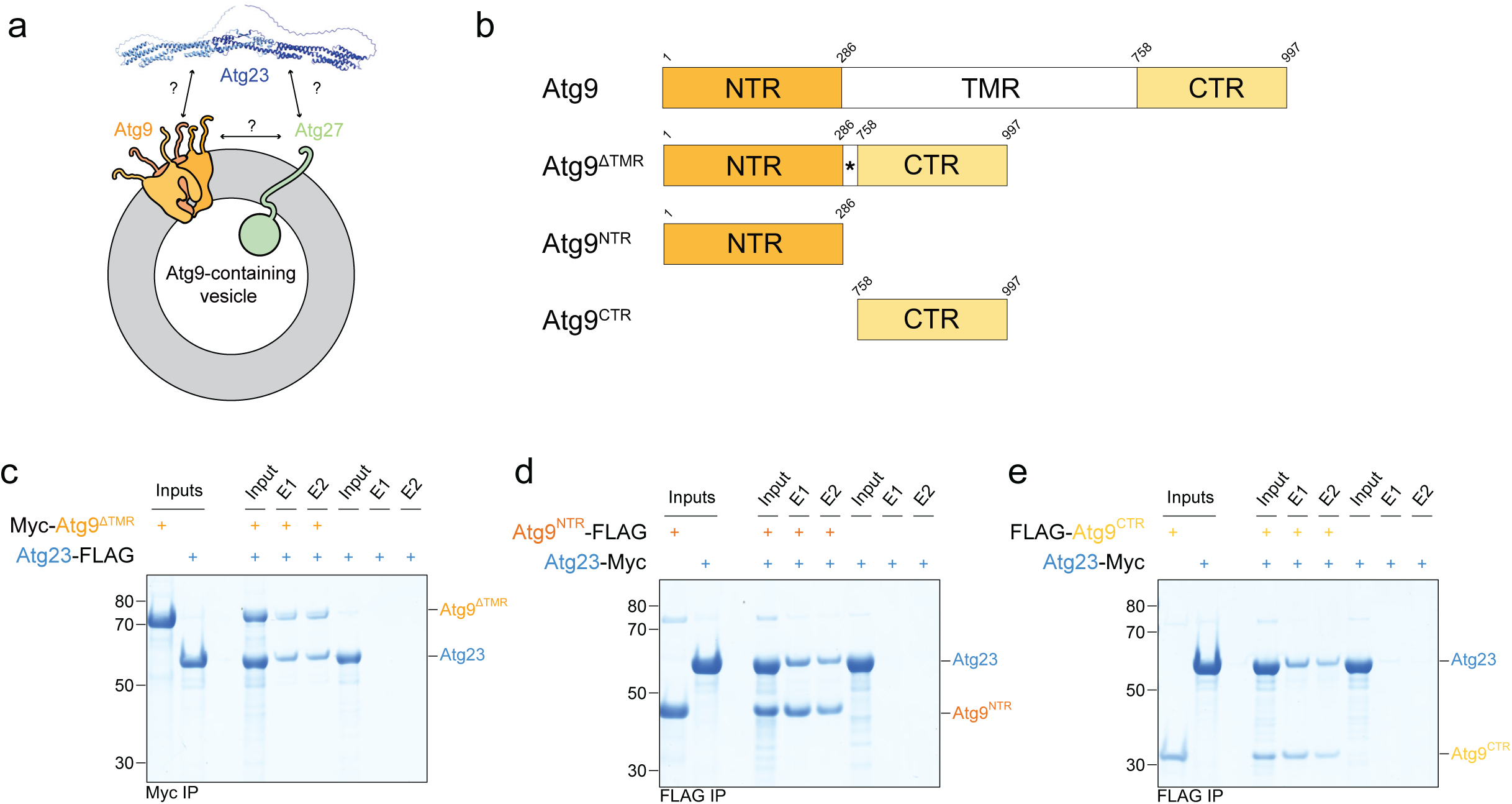
Atg23 interacts with both the N- and C-terminal regions of Atg9. **a**, Schematic representation of an *S. cerevisiae* Atg9-containing vesicle and its core components: Atg9, Atg23 and Atg27. **b**, Schematic domain overview of *S. cerevisiae* Atg9. Atg9 consists of a largely disordered cytoplasmic N- and C-terminal region (NTR and CTR, respectively) and a transmembrane region (TMR). Constructs used in this study (Atg9^ΔTMR^, Atg9^NTR^ and Atg9^CTR^) are shown below. The asterisk refers to a short linker comprised of a small epitope tag (FLAG or Myc). **c**, Atg9 directly interacts with Atg23. Myc-tagged Atg9^ΔTMR^ was immobilized using Myc resin and incubated with FLAG-tagged Atg23. Bound proteins were eluted after washing, and both input and elution (E) samples were analyzed by SDS-PAGE and Coomassie staining. **d-e**, Atg9 interacts with Atg23 via both its N- and C-terminal regions. FLAG-tagged Atg9 constructs, Atg9^NTR^ (panel **d**) and Atg9^CTR^ (panel **e**), were immobilized using FLAG resin and incubated with Myc-tagged Atg23. Following washing, bound proteins were eluted and input and elution (E) fractions were analysed by SDS-PAGE and Coomassie staining.

Using Atg9^ΔTMR^, we next investigated whether Atg9 directly associates with the peripheral membrane protein Atg23. Indeed, when Atg9^ΔTMR^ was immobilized, Atg23 co-eluted in our pulldown experiments, suggesting a direct interaction (**Fig. 1c**). To better understand this interaction, we next tested whether Atg23 interacts with the largely disordered N- and C-terminal regions of Atg9 (Atg9^NTR^ and Atg9^CTR^; **Extended Data Fig. 1c**) and found that both regions were independently capable of binding Atg23 (**Fig. 1d-e**).

### The Atg9^CTR^ interacts with Atg23 via its very C-terminus

We first focused on the interaction between the Atg9^CTR^ and Atg23. Atg23 is thought to form an extended helical homodimer^32^, whose membrane recruitment depends on Atg9^26^. Using mass photometry, we confirmed that Atg23 dimerizes, even at very low nanomolar concentrations (**Extended Data Fig. 2a-b**). We thus used two copies of Atg23 to predict the structure of the complex formed with the Atg9^CTR^ using AlphaFold 3 (AF3)^33^. Consistent with our biochemical analysis, AF3 predicted a complex between the Atg9^CTR^ and the Atg23 dimer with high confidence (**Fig. 2a** and **Extended Data Fig. 2c**). In this model, the extreme C-terminal region of Atg9 interacts with two largely hydrophobic regions in Atg23, which we refer to here as Atg9 binding sites 1 and 2 (BS1 and BS2; **Fig. 2a**). These sites are positioned at the periphery of the Atg23 homodimer, distant from the Atg23 dimerization interface (**Fig. 2a**). At the molecular level, the predicted BS1 interface features multiple hydrophobic contacts between the Atg9^CTR^ and Atg23, along with several intermolecular hydrogen bonds: one between glutamate 947 (E947) in Atg9 and lysine 42 (K42) in Atg23; a second between histidine 958 (H958) in Atg9 and aspartate 43 (D43) in Atg23; a third between tyrosine 16 (Y16) in Atg23 and the backbone carbonyl oxygen of proline 956 (P956) in Atg9; and two additional hydrogen bonds involving aspartate 50 (D50) in Atg23 - one with the side-chain hydroxyl oxygen of serine 948 (S948) and another with the backbone amide hydrogen of phenylalanine 949 (F949) in Atg9 (**Fig. 2b** and **Extended Data Fig. 2d**). In BS2, the extreme C-terminus of Atg9 forms a short *a*-helix that docks into a hydrophobic canyon within a helical bundle in the outward facing tip of Atg23, with a hydrogen bond between lysine 987 (K987) in Atg9 and glutamate 380 (E380) in Atg23 further stabilizing this interaction (**Fig. 2c** and **Extended Data Fig. 2e-f**).

**Fig. 2:**
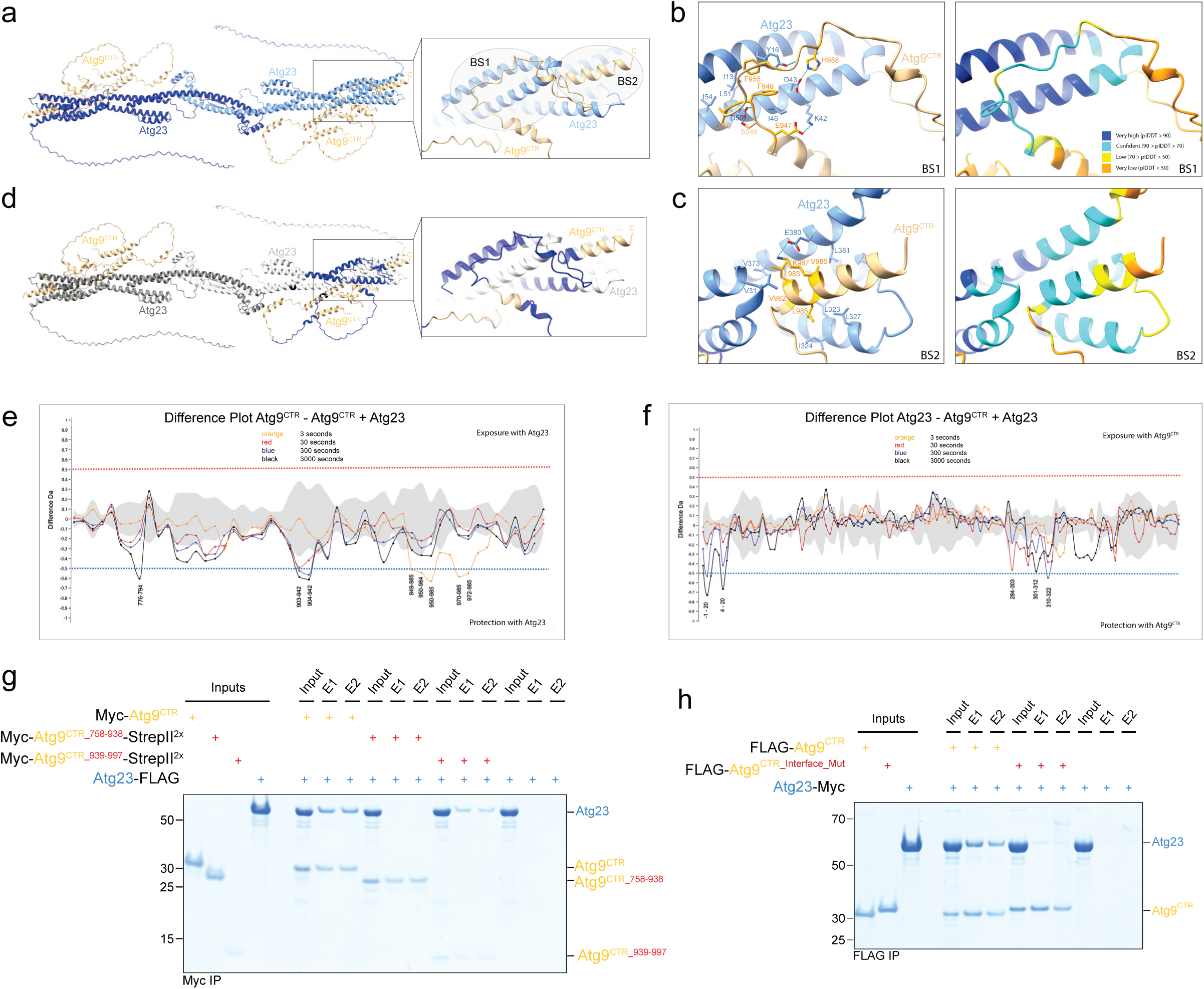
Atg23 interacts with the very C-terminus of Atg9. **a**, AF3 model of the predicted Atg9^CTR^-Atg23 complex showing the interaction interface between the Atg9^CTR^ and Atg23. The Atg9^CTR^ interacts with two different regions in Atg23, referred to as Atg9 binding sites 1 and 2 (BS1 and BS2). **b-c**, AF3 model showing the hydrophobic interface and the hydrogen bonds (cyan dotted lines) stabilizing the Atg9^CTR^-Atg23 interaction in BS1 (panel **b**) and BS2 (panel **c**). The AF3 model is color-coded according to the pIDDT values (right panels). The accuracy of the prediction for regions with pIDDT values ≥ 70 (dark and light blue) is high. Atg9 residues mutated in the Atg9^CTR_Interface_Mut^ mutant are highlighted in orange. **d**, The AF3 model shown in Fig. 2a is colored according to the hydrogen-deuterium exchange (HDX) mass spectrometry (MS) data (shown in Fig. 2e-f). Regions in blue correspond to peptides in the Atg9^CTR^ and Atg23 that are significantly protected from hydrogen-deuterium exchange upon Atg9^CTR^-Atg23 complex formation (HDX MS data is only mapped onto the one protomer shown on the right). Regions in black were not detected in the HDX MS data. **e-f**, HDX MS analysis of the Atg9^CTR^-Atg23 complex. Difference plots comparing the hydrogen-deuterium exchange of the Atg9^CTR^ (panel **e**) and Atg23 (panel **f**) to that of the Atg9^CTR^-Atg23 complex. Negative values below the significance threshold of -0.5 indicate regions that become protected upon complex formation. Deuterium uptake was measured at four time points: 3 seconds (orange), 30 seconds (red), 300 seconds (blue), and 3000 seconds (black). Myc-Atg9^CTR^-StrepII^2x^ and Atg23-Myc were used for the HDX MS analysis with numbering corresponding to the protein sequences without tags. **g**, Atg23 interacts with the very C-terminus of the Atg9^CTR^. Myc-Atg9^CTR^, Myc-Atg9^CTR_758-938^-StrepII^2X^ or Myc-Atg9^CTR_939-997^-StrepII^2x^ were immobilized using Myc resin and incubated with FLAG-tagged Atg23. After washing the resin, bound proteins were eluted and input and elution (E) fractions were analysed by SDS-PAGE and Coomassie staining. **h**, Mutating critical interface residues in the Atg9^CTR^ abolishes the interaction with Atg23. FLAG-tagged Atg9^CTR^ or the Atg9^CTR_Interface_Mut^ mutant _(Atg9CTR_E947A_F949A_L950A_F955A_H958A_V982A_L983A_L985A_V986A_K987A) were immobilized using FLAG resin_ and incubated with Myc-tagged Atg23. After washing the resin, bound proteins were eluted and input and elution (E) fractions were analysed by SDS-PAGE and Coomassie staining.

To verify the AF3 prediction, we performed hydrogen-deuterium exchange mass spectrometry (HDX-MS) experiments, comparing the hydrogen-deuterium exchange in the Atg9^CTR^-Atg23 complex with the exchange in the individual proteins, Atg9^CTR^ and Atg23, respectively. Samples were incubated in deuterated buffer for 3, 30, 300 and 3000 seconds, followed by digestion with pepsin. The resulting peptides were separated by liquid chromatography, and the relative deuterium uptake of all detected peptides was measured by mass spectrometry. Peptides that exhibited reduced deuterium incorporation upon complex formation indicate regions involved in binding or conformational changes^34^. The peptides in Atg23 and the Atg9^CTR^ that were protected from deuterium exchange largely corresponded to the regions predicted by AF3 to form the Atg23-Atg9^CTR^ interface (**Fig. 2d-f**), providing critical experimental support for the AF3 structural model.

To further validate our model biochemically, we expressed and purified an Atg9^CTR^ mutant lacking the region predicted to bind Atg23 (Atg9^CTR_758–938^). As expected, this mutant was unable to bind Atg23 (**Fig. 2g**). Conversely, the extreme C-terminal region of Atg9 (residues 939 to 997; Atg9^CTR_939-997^) was sufficient to mediate Atg23 binding (**Fig. 2g**), further confirming the location of the Atg23 binding site.

To directly validate the interface, we generated an Atg9 mutant in which all residues predicted by the AF3 model to mediate the Atg9^CTR^-Atg23 interaction (excluding those predicted to be required for structural integrity) were substituted with alanine (**Fig. 2b-c**). As expected, the resulting interface mutant, Atg9^CTR_Interface_Mut^ _(Atg9CTR_E947A_F949A_L950A_F955A_H958A_V982A_L983A_L985A_V986A_K987A), exhibited only very weak binding to_ Atg23 (**Fig. 2h**), further confirming the interface. Additionally, we also mutated Atg23 residues predicted to form intermolecular hydrogen bonds in BS1 (Y16, K42 and D43) and BS2 (E380) to alanine (**Extended Data Fig. 2d-e**). However, the resultant Atg23^Y16A_K42A_D43A_E380A^ mutant was still able to bind the Atg9^CTR^ (**Extended Data Fig. 2g**), suggesting that complex formation is primarily driven by hydrophobic interactions.

### The Atg9^NTR^ interacts with Atg23 via its extreme N-terminus, occupying the same binding site as the Atg9^CTR^

Our experiments clearly demonstrate that it is not only the Atg9^CTR^ that interacts with Atg23 but also the Atg9^NTR^ (**Fig. 1d-e**). This bipartite binding mode also explains why Atg23 binding was unchanged for an Atg9^ΔTMR^ mutant with the experimentally verified Atg23 binding site in the Atg9 C-terminus deleted (Atg9^ΔTMR_Δ939-997^) (**Extended Data Fig. 3a**). To dissect this additional binding site in more detail, we next sought to better understand the Atg9^NTR^-Atg23 interaction.

The Atg9 N-terminus is predicted to be largely unstructured^35^ (**Extended Data Fig. 1c**). Using two copies of Atg23 and a single copy of the Atg9^NTR^ as input sequences, AF3 identified two potential Atg23 binding sites within the Atg9^NTR^: one in the very N-terminus and another closer to its C-terminus (**Fig. 3a**). As the AF3 prediction showed only intermediate confidence, we performed additional HDX-MS experiments to further investigate the Atg9^NTR^-Atg23 interface. However, the only region with weak but significant protection from deuterium exchange was a C-terminal segment within the Atg9^NTR^ (**Extended Data Fig. 3b**), which differed from the two regions implicated in complex formation by the AF3 model (**Fig. 3a**).

**Fig. 3:**
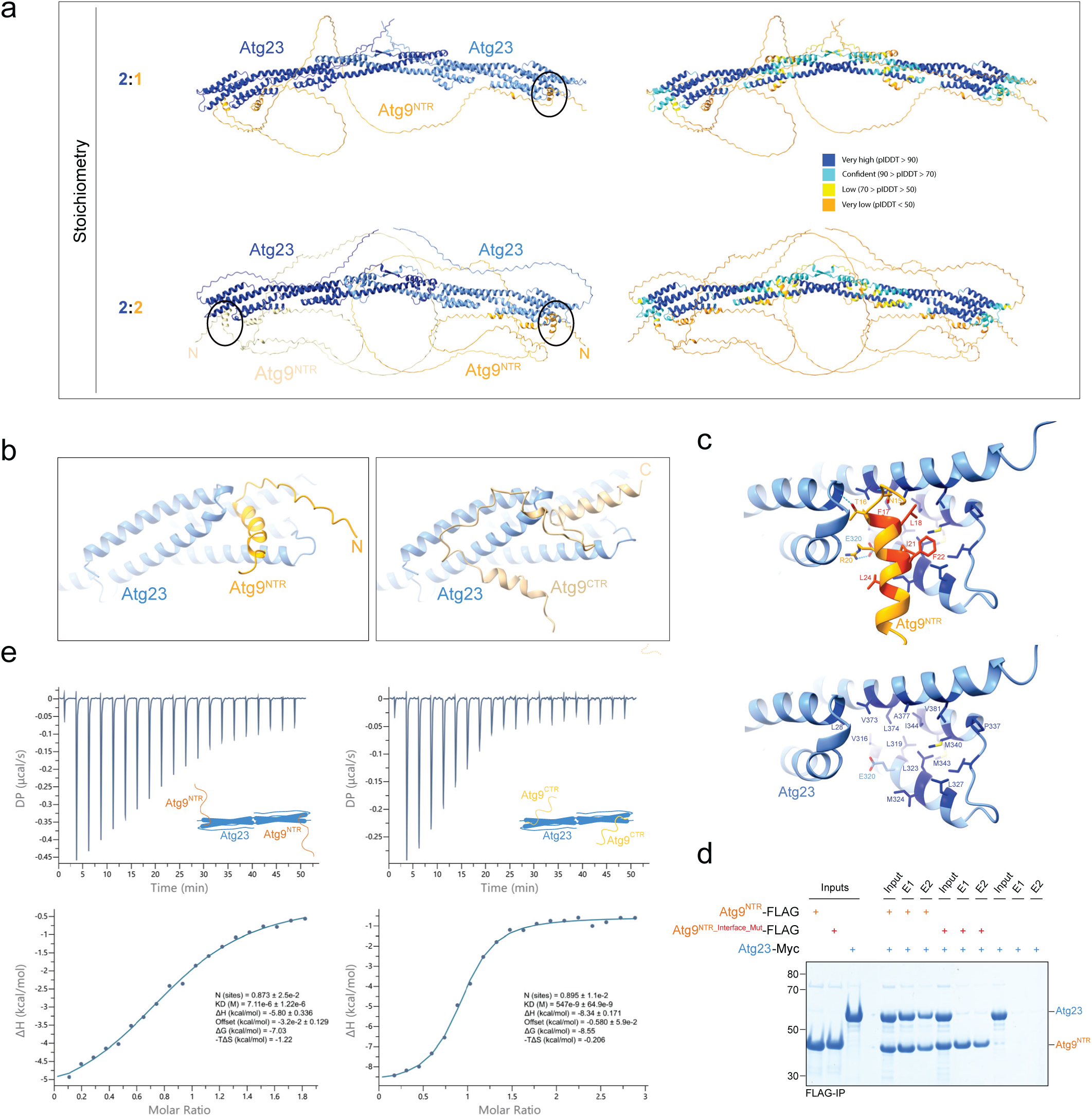
Atg23 also interacts with the Atg9^NTR^, using the same binding site it employs for binding the Atg9^CTR^. **a**, AF3 predictions of the Atg9^NTR^-Atg23 complex. Two different stoichiometries of the Atg23-Atg9^NTR^ complex were modeled and compared: 2:1 (upper panel) and 2:2 (lower panel). Atg23 was included as a homodimer in both cases. AF3 predictions indicate that the extreme N-terminus of the Atg9^NTR^ interacts with Atg23 (indicated by black circles). Model confidence is visualized using pIDDT color-coding (right panels). **b**, The Atg9^NTR^ and Atg9^CTR^ binding sites on Atg23 partially overlap. AF3 models of the Atg9^NTR^-Atg23 (left) and Atg9^CTR^-Atg23 (right) complexes were aligned to compare Atg9^NTR^ and Atg9^CTR^ binding to Atg23. **c**, AF3 model of the Atg9^NTR^-Atg23 interface. Stabilizing intermolecular hydrogen bonds are depicted as cyan dotted lines and hydrophobic interface residues in Atg23 and Atg9 are shown in dark blue and red, respectively (upper panel). The Atg9 residues in red were mutated to alanine in the Atg9^NTR_Interface_Mut^ mutant. The largely hydrophobic Atg9^NTR^ binding site on Atg23 is depicted without the Atg9^NTR^ in the lower panel. **d**, Biochemical validation of the predicted Atg9^NTR^-Atg23 interface. The FLAG-tagged wild-type and Atg9^NTR^ interface mutant (Atg9^NTR_Interface_Mut^: Atg9^NTR_F17A_L18A_I21A_F22A_L24A^) were immobilized using FLAG resin and incubated with Myc-tagged Atg23. After washing the resin, bound proteins were eluted and input and elution (E) fractions were analysed by SDS-PAGE and Coomassie staining. **e**, Isothermal titration calorimetry (ITC) analysis of the Atg9^NTR^-Atg23 (left) and Atg9^CTR^-Atg23 (right) interaction. Raw heat changes (DP, µcal/s) (top) and integrated binding isotherms (bottom) are shown. Binding isotherms were derived from integrated heat signals and fitted to a One Set of Sites model to derive thermodynamic parameters. FLAG-tagged Atg9^NTR^, FLAG-tagged Atg9^CTR^ and untagged Atg23 were used for the experiments.

To further refine our structural model, we reassessed the prediction using a stoichiometry of 2:2 (**Fig. 3a**). Notably, when two Atg9^NTR^ molecules were included, the confidence of the AF3 model significantly increased (**Fig. 3a** and **Extended Data Fig. 3c**). In this model, Atg23 binds both molecules of the Atg9^NTR^, with the very N-terminus of the Atg9^NTR^ forming a short *a*-helix (amino acids 11-26), that binds a peripheral helical bundle in Atg23 away from the dimerization interface (**Fig. 3a**). Interestingly, the Atg9^NTR^-interacting region on Atg23 overlaps with the BS2 for the Atg9^CTR^ (**Fig. 3b**). However, while the N-terminal *a*-helix of the Atg9^NTR^ is predicted to bind Atg23 in a nearly perpendicular manner, the extreme C-terminal *a*-helix of the Atg9^CTR^ docks into the Atg23 helical bundle in an almost parallel fashion (**Fig. 3b**). Like the Atg9^CTR^-Atg23 interaction, the Atg9^NTR^-Atg23 interface is largely hydrophobic, with Atg9 residues phenylalanine 17 (F17), leucine 18 (L18), isoleucine 21 (I21), phenylalanine 22 (F22) and leucine 24 (L24) docking into a hydrophobic pocket at the distal end of each Atg23 protomer (**Fig. 3c** and **Extended Data Fig. 3d**). Furthermore, the AF3 model suggests that the Atg23-Atg9^NTR^ interaction is further stabilized by three hydrogen bonds: one between arginine 20 (R20) in Atg9 and glutamate 320 (E320) in Atg23; another between threonine 16 (T16) in Atg9 and the backbone carbonyl oxygen of leucine 28 (L28) in Atg23; and a third between asparagine 15 (N15) in Atg9 and the backbone carbonyl oxygen of valine 373 (V373) in Atg23 (**Fig. 3c**).

To validate the AF3 model, we mutated the hydrophobic interface residues in the Atg9^NTR^ to alanine (F17A, L18A, I21A, F22A and L24A) and tested the resultant mutant (Atg9^NTR_Interface_Mut^) in Atg23 binding assays. Consistent with our model, this mutant had almost completely lost its ability to interact with Atg23 (**Fig. 3d**).

Based on the experimentally validated AF3 models, the Atg9^NTR^ and Atg9^CTR^ appear to have overlapping binding sites on Atg23 (**Fig. 3b**). To further investigate this, we compared the binding affinities of Atg23 for the Atg9^NTR^ and Atg9^CTR^ using isothermal titration calorimetry (ITC).The results revealed that Atg23 has an approximately 10-fold higher affinity for the Atg9^CTR^ than for the Atg9^NTR^, with a dissociation constant (K_D_) of 540 ± 80 nM for the Atg23-Atg9^CTR^ interaction and a K_D_ of 6.7 ± 1.2 µM for the Atg23-Atg9^NTR^ interaction (**Fig. 3e**). This difference in binding affinity is likely due to a larger interface between Atg23 and the Atg9^CTR^, as suggested by the AF3 model (**Fig. 2a-c**). Moreover, the n-values for both interactions are approximately 1, suggesting that a single Atg23 dimer binds two Atg9^NTR^ or Atg9^CTR^ molecules (**Fig. 3e**).

These ITC results suggest that the Atg9^CTR^ can outcompete the Atg9^NTR^ for binding to Atg23. To test this experimentally, we incubated the Atg9^NTR^ with Atg23 in the presence or absence of the wild-type Atg9^CTR^ or the Atg9^CTR_Interface_Mut^ mutant. Consistent with the ITC data and our model, the Atg9^NTR^-Atg23 interaction was markedly weakened in the presence of the wild-type Atg9^CTR^, but not in the presence of the Atg9^CTR^ interface mutant (**Extended Data Fig. 3e**). These findings reinforce the predicted competitive binding mode and confirm that the Atg9^CTR^ can outcompete the Atg9^NTR^ for Atg23 binding, supporting a model in which Atg23 preferentially associates with the C-terminal region of Atg9 under steady-state conditions.

### Atg1-mediated phosphorylation of the Atg9^NTR^ and Atg9^CTR^ negatively regulates Atg23 binding

Atg9 has previously been shown to be phosphorylated by the Atg1 kinase^22,23^ and its homologs Ulk1/Ulk2 in higher eukaryotes^36,37^. To investigate how Atg1-mediated phosphorylation influences Atg9’s interaction with Atg23, we performed pulldown experiments after phosphorylating the Atg9^NTR^, the Atg9^CTR^ and Atg23 using recombinant Atg1. Notably, Atg1-mediated phosphorylation destabilized the Atg9^CTR^-Atg23 interaction (**Fig. 4a**), and completely abolished the Atg9^NTR^-Atg23 interaction (**Fig. 4b**). To determine whether these effects were due to the phosphorylation of Atg23 or Atg9, we individually phosphorylated Atg23, the Atg9^NTR^, or the Atg9^CTR^, then depleted ATP using apyrase prior to adding the non-phosphorylated binding partner. Atg1-mediated phosphorylation of Atg23 did not significantly affect its interaction with Atg9^NTR^ and Atg9^CTR^; however, phosphorylation of both Atg9^NTR^ and Atg9^CTR^ interfered with Atg23 binding, fully mimicking the effect observed when both binding partners were phosphorylated by Atg1 (**Fig. 4a**-**b**). These results demonstrate that it is phosphorylation of Atg9 and not Atg23 that regulates the Atg9-Atg23 interaction.

**Fig. 4:**
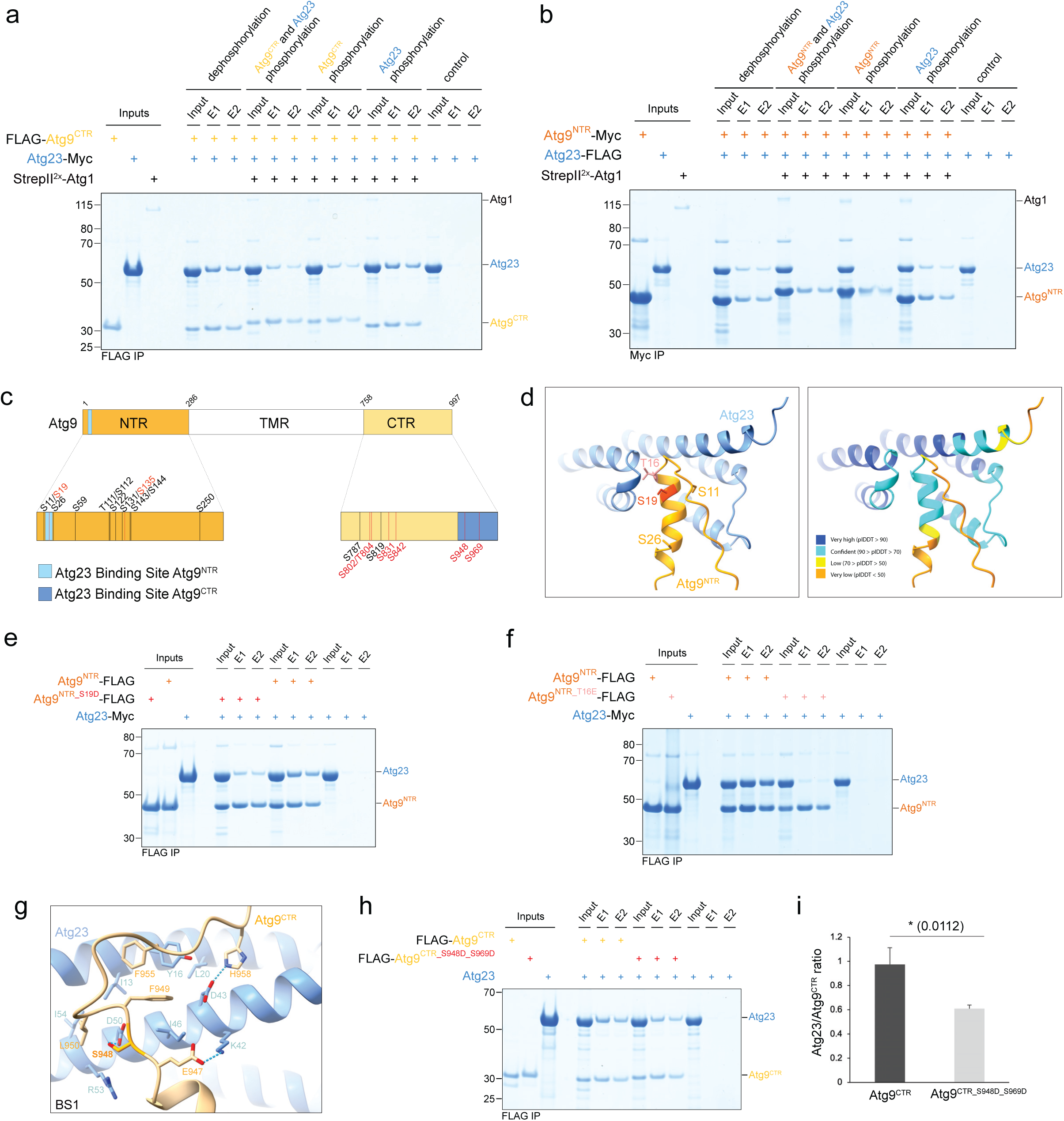
The Atg9-Atg23 interaction is inhibited by Atg1-mediated phosphorylation of Atg9. **a-b**, Atg1-dependent phosphorylation of the Atg9^NTR^ and Atg9^CTR^ regulates Atg23 binding. FLAG-tagged Atg9^CTR^ (panel **a**) or Myc-tagged Atg9^NTR^ (panel **b**) and Atg23 were phosphorylated using substoichiometric amounts of recombinant Atg1. ATP was depleted using apyrase prior to combining the indicated phosphorylated and non-phosphorylated proteins. As a control, the Atg9^NTR^ or Atg9^CTR^, and Atg23 were dephosphorylated using - phosphatase (-PP). The Atg9^CTR^ or Atg9^NTR^ were immobilized using FLAG or Myc resin (panel **a** and **b**, respectively) and following washing, bound proteins were eluted. Both input and elution (E) samples were analyzed by SDS-PAGE and Coomassie staining. **c**, Domain overview of *S. cerevisiae* Atg9. Previously reported phosphorylation sites in the Atg9^NTR^ and Atg9^CTR^ are indicated. All Atg1-dependent phosphorylation sites are highlighted in red and the Atg23 binding sites in the Atg9 NTR and CTR in light and dark blue, respectively. **d**, Mapping of phosphorylation sites onto the Atg9^NTR^-Atg23 interface. Previously reported phosphorylation sites within the Atg9^NTR^-Atg23 binding region and the potential phosphorylation site T16 are mapped onto the AF3-predicted structure^23,38,51^. Serine 19 (S19), which is phosphorylated by Atg1 *in vivo*^23^, is also indicated. The AF3 model is color-coded according to the pIDDT values (right), with high-confidence regions (pIDDT value ≥ 70) shown in dark and light blue. **e-f**, Phosphomimicking mutations of S19 and T16 interfere with Atg23 binding. FLAG-tagged Atg9^NTR^ and the phosphomimicking mutants Atg9^NTR_S19D^ (panel **e**) and Atg9^NTR_T16E^ (panel **f**) were incubated with Myc-tagged Atg23. Proteins were immobilized on FLAG resin, washed, and eluted. Input and elution (E) samples were analyzed by SDS-PAGE and Coomassie staining. **g**, Serine 948 (S948) localizes to the predicted Atg9^CTR^-Atg23 interface. In the AF3-predicted model, S948 forms a hydrogen bond with aspartate 50 (D50) of Atg23 at Atg9 binding site 1 (BS1), providing a possible structural explanation for how Atg1-mediated phosphorylation at this site reduces Atg23 binding. **h**, Phosphomimicking mutations of S948 and S969 reduce Atg23 binding. FLAG-tagged Atg9^CTR^ and the phosphomimicking mutant Atg9^CTR_S948D_S969D^ were incubated with Atg23. Proteins were immobilized on FLAG resin. After washing the resin, bound proteins were eluted and input and elution (E) fractions were analysed by SDS-PAGE and Coomassie staining. **i**, Quantification of pulldown experiments shown in panel **h**). Sypro Ruby-stained gels were used to quantify Atg23 binding (n = 3). The mean Atg23/Atg9^CTR^ ratio was calculated from the E1 fractions of three independent pulldown assays using FLAG-tagged Atg9^CTR^ or Atg9^CTR_S948D_S969D^ as bait. Error bars represent standard deviation. An unpaired t-test revealed a significant difference between the Atg23/Atg9^CTR^ ratios in the presence of Atg9^CTR^ and Atg9^CTR_S948D_S969D^ (t = 4.4509, *P* = 0.0112).

To understand the molecular basis of this posttranslational regulation, we first focused on the Atg9^NTR^-Atg23 interaction and the previously reported phosphorylation sites within the Atg9^NTR^-Atg23 interface^22,23,38^ (**Fig. 4c**). Notably, serine 19 (S19), previously reported to be phosphorylated by Atg1^23^, mapped exactly to the Atg9^NTR^-Atg23 interface (**Fig. 4c**-**d**). We thus mutated S19 to aspartate to mimic its phosphorylation and tested the resultant mutant, Atg9^NTR_S19D^, in pulldown experiments. Strikingly, the mutation significantly interfered with Atg23 binding (**Fig. 4e**), suggesting that Atg1-mediated phosphorylation of S19 regulates the Atg9-Atg23 interaction.

In addition to S19, the AF3 model suggested that phosphorylation of threonine 16 (T16) may also regulate Atg23 binding (**Fig. 4d**). Although T16 has not previously been identified as a phosphorylation site, it may have been missed due to the short length of its tryptic peptide. At just six amino acids, it falls below the typical seven-amino-acid threshold commonly used in standard peptide identification workflows. The positioning of T16 relative to Atg23 suggests that its phosphorylation could sterically interfere with complex formation (**Fig. 4d**). Moreover, its surrounding sequence complies with the previously published Atg1 consensus motif^22^. Therefore, we also tested the corresponding phosphomimicking mutant, Atg9^NTR_T16E^, and observed that Atg23 binding was severely impaired (**Fig. 4f**), mimicking the effect of the Atg9^NTR_Interface_Mut^ interface mutant.

Given that serine 11 (S11) and serine 26 (S26) have also been reported as phosphorylation sites^22,23,38^ and due to their proximity to T16 and S19 (**Fig. 4d**), we also tested the corresponding phosphomimicking mutants, Atg9^NTR_S11D^ and Atg9^NTR_S26D^, in pulldown assays. However, consistent with their position in the AF3 model, neither mutation significantly influenced Atg23 binding (**Extended Data Fig. 4a-b**).

Finally, to show that it is only phosphorylation of S19 and possibly T16 that regulates Atg23 binding, we also investigated whether phosphorylation outside of the very N-terminus influences Atg23 binding. We thus generated and analysed two phosphomimicking _mutants, Atg9NTR_T111E_S112D_S122D_S131D_S135D_S143D_S144D and Atg9NTR_S209D_S210D_S220D_S241D_S250D, in_ which all serine and threonine residues in either the central or C-terminal region of the Atg9^NTR^ were replaced by aspartate or glutamate residues, respectively (**Extended Data Fig. 4c-d**). Neither of these mutants had a significantly reduced affinity for Atg23, further confirming that it is phosphorylation of the very N-terminus that regulates the interaction of the Atg9^NTR^ with Atg23.

As Atg1-mediated phosphorylation also impacts the Atg9^CTR^-Atg23 interaction (**Fig. 4a**), we next investigated which Atg1-dependent phosphorylation sites regulate this interaction. Like the Atg9^NTR^, the Atg9^CTR^ has been shown to be extensively phosphorylated, with most phosphorylation sites being Atg1 targets^22–24,38^ (**Fig. 4c**). Atg1 is a serine/threonine kinase^23^. We therefore focused on Atg9 BS1 (**Fig. 2a-b**), as none of the interface residues in BS2 are serines or threonines and mutating the two tyrosines, Y989 and Y990, to non-phosphorylatable phenylalanines did not interfere with the Atg1-dependent destabilization of the Atg9^CTR^-Atg23 interaction (**Extended Data Fig. 4e**). Serine 948 (S948) and serine 969 (S969) had both previously been shown to be phosphorylated by Atg1^22–24^ (**Fig. 4c**). While S969 is not directly in contact with Atg23 according to the AF3 model (**Extended Data Fig. 4f**), S948 maps to a highly kinked segment in the Atg9^CTR^ that interacts with Atg23 in BS1 and has a high prediction confidence (**Fig. 4g** and **Extended Data Fig. 4f**). To test whether phosphorylation of S948 and S969 causes the destabilization of the Atg9^CTR^-Atg23 interaction, we mutated both residues in the Atg9^CTR^ to alanine. For the resultant mutant, Atg9^CTR_S948A_S969A^, Atg23 binding was no longer sensitive to Atg1-mediated phosphorylation (**Extended Data Fig. 4g**), confirming identification of the causative phosphorylation sites.

Consistently, the corresponding phosphomimicking mutant, Atg9^CTR_S948D_S969D^, showed significantly reduced Atg23 binding (**Fig. 4h-i**), recapitulating the effect observed for Atg1-mediated phosphorylation. Based on the AF3 model, phosphorylation of S948 likely interferes with hydrogen bond formation resulting in the electrostatic repulsion with aspartate 50 (D50) in Atg23 (**Fig. 2b** and **Extended Data Fig. 4f**). Notably, arginine, R53, is adjacent to D50 in Atg23 (**Fig. 4g**), which may mitigate this repulsion. However, this could lead to the repositioning of phenylalanines 949 (F949) and 955 (F955) away from tyrosine 16 (Y16) in Atg23, disrupting their predicted u-u stacking interactions and weakening the Atg23-Atg9^CTR^ interaction in BS1. As a result, only the extreme C-terminus of Atg9 likely remains engaged with Atg23 in BS2, explaining the reduced affinity upon Atg1-mediated phosphorylation.

### Atg23 needs to interact with both the Atg9^NTR^ and Atg9^CTR^ to support autophagy

Having characterised the Atg9-Atg23 interaction and its phosphoregulation *in vitro*, we next sought to confirm the physiological relevance of the two Atg23 binding sites in Atg9. We therefore performed Pho8Δ60 assays^39^ to assess bulk autophagy in yeast expressing either the Atg9^NTR_Interface_Mut^ or Atg9^CTR_Interface_Mut^ mutant. While bulk autophagy was significantly reduced in yeast expressing the Atg9^CTR_Interface_Mut^ mutant, those expressing the Atg9^NTR_Interface_Mut^ mutant exhibited a markedly more severe defect (**Fig. 5a**). This observation was unexpected, as the Atg9^CTR^ displays approximately tenfold higher affinity for Atg23 compared to the Atg9^NTR^ (**Fig. 3e**). We therefore aimed to validate the effect of disrupting the Atg9^NTR^-Atg23 interaction using an alternative strategy. *In vitro*, the phosphomimicking T16E and S19D mutations in Atg9 also interfered with Atg23 binding, with the T16E mutation alone effectively abrogating Atg23 binding (**Fig. 4e-f**). Although less pronounced than the autophagy defect observed for the Atg9^NTR_Interface_Mut^ mutant, yeast expressing the Atg9^S19D^, Atg9^T16E^ or the Atg9^T16E_S19D^ double mutant also exhibited a significant autophagy defect in Pho8Δ60 assays (**Fig. 5b**). Together, these results show that both the Atg9^NTR^-Atg23 and the Atg9^CTR^-Atg23 interactions play a role in bulk autophagy, as either interaction alone can only partially compensate for the loss of the other.

**Fig. 5:**
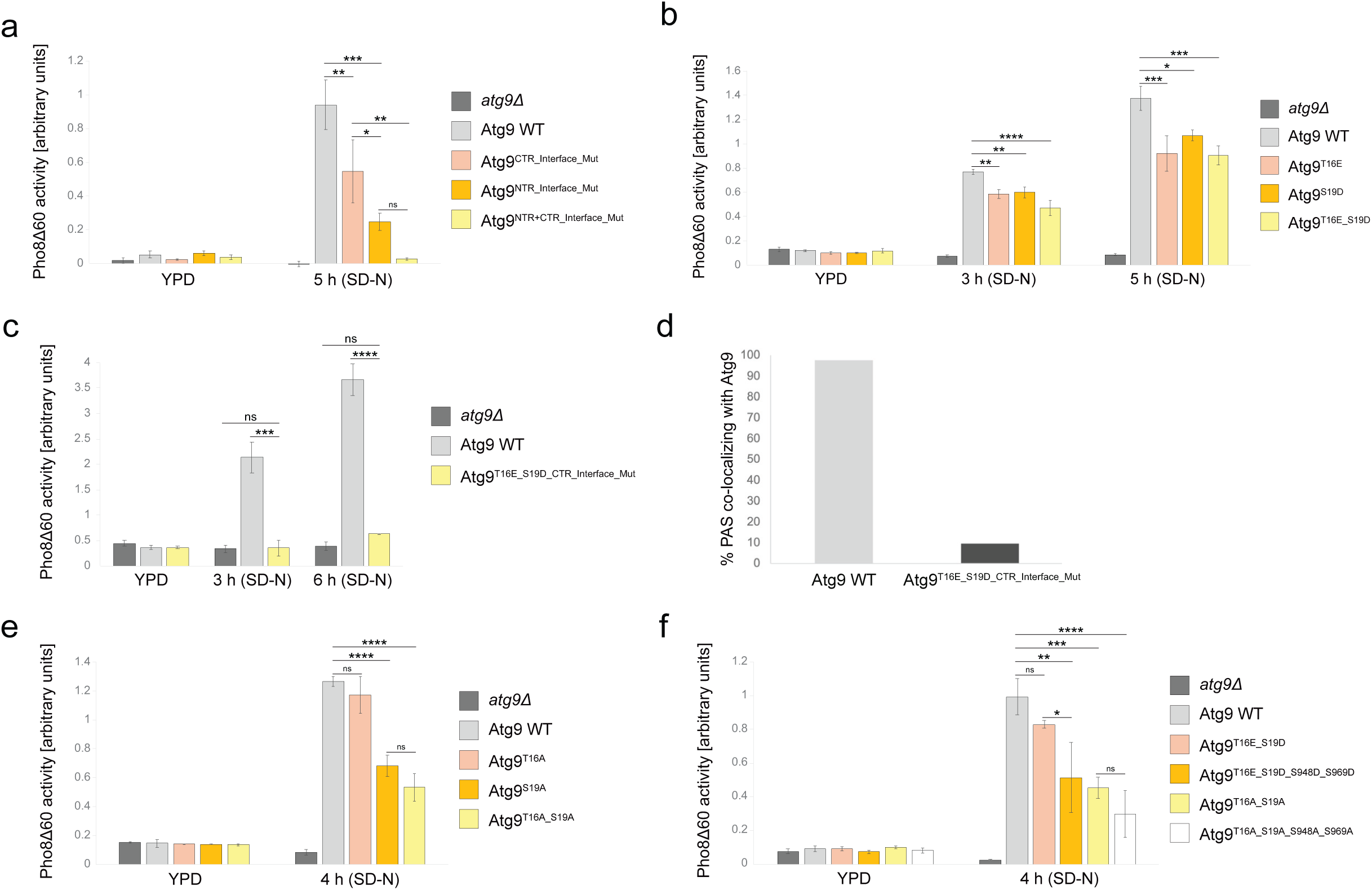
The Atg9-Atg23 interaction and its regulation by Atg1 are required for bulk autophagy. **a-b**, Both the Atg9^NTR^-Atg23 and Atg9^CTR^-Atg23 interactions are required for bulk autophagy. Panel **a** shows Pho8Δ60 assay results measuring bulk autophagy in *S. cerevisiae* strains expressing either wild-type Atg9, an N-terminal interface mutant (Atg9^NTR_Interface_Mut^: _Atg9F17A_L18A_I21A_F22A_L24A), a C-terminal interface mutant (Atg9CTR_Interface_Mut: Atg9E947A_F949A_L950A_F955A_H958A_V982A_L983A_L985A_V986A_K987A) or a mutant combining the N- and C-_ terminal interface mutations (Atg9^NTR+CTR_Interface_Mut^). Panel **b** shows Pho8Δ60 assay results for *S. cerevisiae* strains expressing wild-type Atg9 or Atg9 mutants containing phosphomimicking substitutions in the NTR (Atg9^T16E^, Atg9^S19D^ and Atg9^T16E_S19D^). An *atg9Δ* strain was included as a negative control. Cells were grown exponentially in YPD medium or subjected to nitrogen-starvation (SD-N) for 5 (panel **a**) or 3 and 5 (panel **b**) hours. Pho8Δ60-specific alkaline phosphatase activity was quantified, with mean values from three biological replicates (n = 3) presented. Error bars indicate the standard deviation. Statistical significance between yeast strains was assessed using one-way ANOVA followed by Tukey’s multiple comparisons test. **c**, Disrupting both Atg23 binding sites in Atg9 impairs bulk autophagy. Pho8Δ60 experiments were carried out and analysed as described in Fig. 5a-b. A budding yeast strain expressing Atg9 with mutations that simultaneously disrupt both the N- and C-terminal interfaces with Atg23 (Atg9^T16E_S19D_CTR_Interface_Mut^: _Atg9T16E_S19D_E947A_F949A_L950A_F955A_H958A_V982A_L983A_L985A_V986A_K987A) was compared to cells_ expressing wild-type Atg9 and to an *atg9Δ* strain. Cells were grown exponentially in YPD medium or subjected to nitrogen-starvation (SD-N) for 3 and 6 hours. **d**, The Atg9^T16E_S19D_CTR_Interface_Mut^ mutant exhibits strongly impaired PAS localization. Quantification of fluorescence microscopy experiments (see Extended Data Fig. 5d) of cells expressing Atg1^D211A^-mTagBFP2 as a PAS marker and either EGFP-tagged wild-type Atg9 or the Atg9^T16E_S19D_CTR_Interface_Mut^ mutant. Cells were grown exponentially in nutrient-rich YPD medium, and the percentage of PAS puncta with co-localizing Atg9 signal was quantified and plotted. **e-f**, Atg1-mediated phosphoregulation of the Atg9-Atg23 interaction seems largely driven by phosphorylation of the Atg9^NTR^, particularly at S19. Pho8Δ60 experiments were carried out and analysed as described in Fig. 5a-b. Budding yeast expressing various Atg9 mutants with different non-phosphorylatable alanine substitutions in the NTR (Atg9^T16A^, Atg9^S19A^ and Atg9^T16A_S19A^; panel **e**) or non-phosphorylatable or phosphomimicking mutations in only the NTR or in the NTR and CTR (Atg9^T16A_S19A^, Atg9^T16E_S19D^, Atg9^T16A_S19A_S948A_S969A^ and Atg9^T16E_S19D_S948D_S969D^; panel **f**) were compared to cells expressing wild-type Atg9 and an *atg9Δ* strain. Cells were grown exponentially in YPD medium or subjected to nitrogen-starvation (SD-N) for 4 hours. All Atg9 variants tested contained a C-terminal 3xFLAG tag (panels **a-c** and **e-f**).

The sharp contrast between the phenotypes of the phosphomimicking T16E and S19D mutants and the Atg9^NTR_Interface_Mut^ mutant (**Fig. 5a-b**) suggests distinct underlying mechanisms. This interpretation is supported by our *in vitro* and *in vivo* evidence that all Atg9^NTR^ variants are properly folded, as indicated by their ability to bind Atg11 (**Extended Data Fig. 5a**), and that their expression levels are comparable to wild-type Atg9 (**Extended Data Fig. 5b**). These findings suggest that at least one of the five residues mutated in the Atg9^NTR_Interface_Mut^ mutant plays additional roles in autophagy, possibly by mediating interactions with other, as-yet unidentified proteins.

Thus, to determine the real impact of simultaneously disrupting both the N- and C-terminal Atg23-binding sites in Atg9, we generated two Atg9 mutants by combining the C-terminal interface mutations with either S19D or both T16E and S19D. Strikingly, the _resultant Atg9S19D_CTR_Interface_Mut and Atg9T16E_S19D_CTR_Interface_Mut mutants exhibited a severe bulk_ autophagy defect, with the Atg9^T16E_S19D_CTR_Interface_Mut^ mutant behaving similarly to an *atg9Δ* strain (**Fig. 5c** and **Extended Data Fig. 5c**), suggesting that the Atg9-Atg23 interaction is essential for bulk autophagy.

Consistent with the physiological importance of Atg23 in Atg9-containing vesicle formation^6^, fluorescence microscopy analysis revealed that the Atg9^T16E_S19D_CTR_Interface_Mut^ mutant failed to localize to the site of autophagosome formation in cells expressing a catalytically inactive, mTagBFP2-tagged Atg1 mutant (Atg1^D211A^-mTagBFP2) as a PAS marker (**Fig. 5d** and **Extended Data Fig. 5d**). This observation aligns with the severe autophagy defect of the same mutant in Pho8Δ60 assays (**Fig. 5c**), further underscoring the critical role of the Atg9-Atg23 interaction in autophagy.

### Atg1-mediated phosphoregulation of the Atg9-Atg23 interaction is critical for autophagy

Having established the role of the Atg9-Atg23 interaction in autophagy, we next sought to assess the physiological relevance of its phosphoregulation. We therefore expressed different non-phosphorylatable alanine mutants of Atg9 in *S. cerevisiae* and performed Pho8Δ60 assays to evaluate their impact on bulk autophagy. We first examined phosphorylation sites that regulate the Atg9^NTR^-Atg23 interaction by expressing the Atg9^S19A^ mutant. To confirm that the S19A mutation alters only the phosphorylation status of Atg9 without affecting Atg23 binding, we performed pulldown assays using the Atg9^NTR_S19A^ mutant and Atg23. Binding of Atg23 was unaffected (**Extended Data Fig. 5e**), validating the Atg9^NTR_S19A^ mutant as a suitable tool to study the phosphorylation-dependent regulation of the Atg9^NTR^-Atg23 interaction *in vivo*. Strikingly, the Atg9^S19A^ mutant exhibited a strong reduction in bulk autophagy upon nitrogen starvation (**Fig. 5e**), indicating that Atg1-mediated phosphorylation of S19 plays a key role in autophagy, at least in part by regulating Atg23 binding.

To assess whether phosphorylation of T16 also contributes to autophagy regulation, we measured bulk autophagy in yeast expressing the Atg9^T16A^ mutant. Expression of this non-phosphorylatable mutant did not result in a significant bulk autophagy defect (**Fig. 5e** and **Extended Data Fig. 5f**), suggesting that phosphorylation of T16 does not play a major role in autophagy regulation. Consistently, combining the T16A and S19A mutations did not produce a more severe phenotype than the Atg9^S19A^ mutant (**Fig. 5e**), confirming that S19 phosphorylation alone is sufficient to regulate the Atg9^NTR^-Atg23 interaction.

To gain further insights into the phosphoregulation of the Atg9-Atg23 interaction and to evaluate the contribution of C-terminal phosphorylation sites in Atg9 (S948 and S969), we combined N- and C-terminal phosphomimicking or alanine mutations and assessed bulk autophagy in the resultant Atg9 mutants. The phosphorylation-deficient Atg9^T16A_S19A_S948A_S969A^ mutant exhibited only a mild, non-significant additional reduction in bulk autophagy compared to the Atg9^T16A_S19A^ mutant (**Fig. 5f**). By contrast, the phosphomimicking Atg9^T16E_S19D_S948D_S969D^ mutant showed a significantly stronger autophagy defect than the Atg9^T16E_S19D^ mutant (**Fig. 5f**), which likely reflects an additive effect caused by the simultaneous disruption of Atg23 binding to both the Atg9^NTR^ and Atg9^CTR^ regions. As S19, S948 and S969 are all phosphorylated by Atg1 *in vivo*^22–24^, these findings suggest that phosphorylation at these sites modulates autophagy by destabilizing the Atg9-Atg23 interaction at the PAS. The minimal additional defect observed in the Atg9^T16A_S19A_S948A_S969A^ mutant compared to Atg9^T16A_S19A^ indicates that this regulatory mechanism is largely driven by Atg1-mediated phosphorylation of the Atg9 N-terminus, particularly at S19 (**Fig. 5f**).

### Atg27 and Atg9 directly interact via their C-terminal cytoplasmic regions

Atg9-containing vesicles not only contain Atg9 but also the type I transmembrane protein Atg27^25^, which contains an N-terminal luminal mannose 6-phosphate receptor homology (MRH) domain, a single transmembrane helix and a disordered C-terminal tail facing the cytoplasm^27^ (**Fig. 6a**). To better understand the function of Atg27 in autophagy, we wanted to first investigate whether Atg27 directly interacts with Atg9 or Atg23. Attempts to co-express and purify full-length Atg9 and Atg27 to study complex formation were inconclusive, as the high detergent concentrations required during membrane solubilization likely interfered with complex stability. As the luminal domain of Atg27 is not required for autophagy (**Extended Data Fig. 6a**) and Atg9 has no significant luminal domains or loops^14,16–18^, we focused on the C-terminal cytoplasmic region of Atg27 (Atg27^CTR^: amino acids 223-271) to understand its interactions with the remaining vesicle components.

**Fig. 6:**
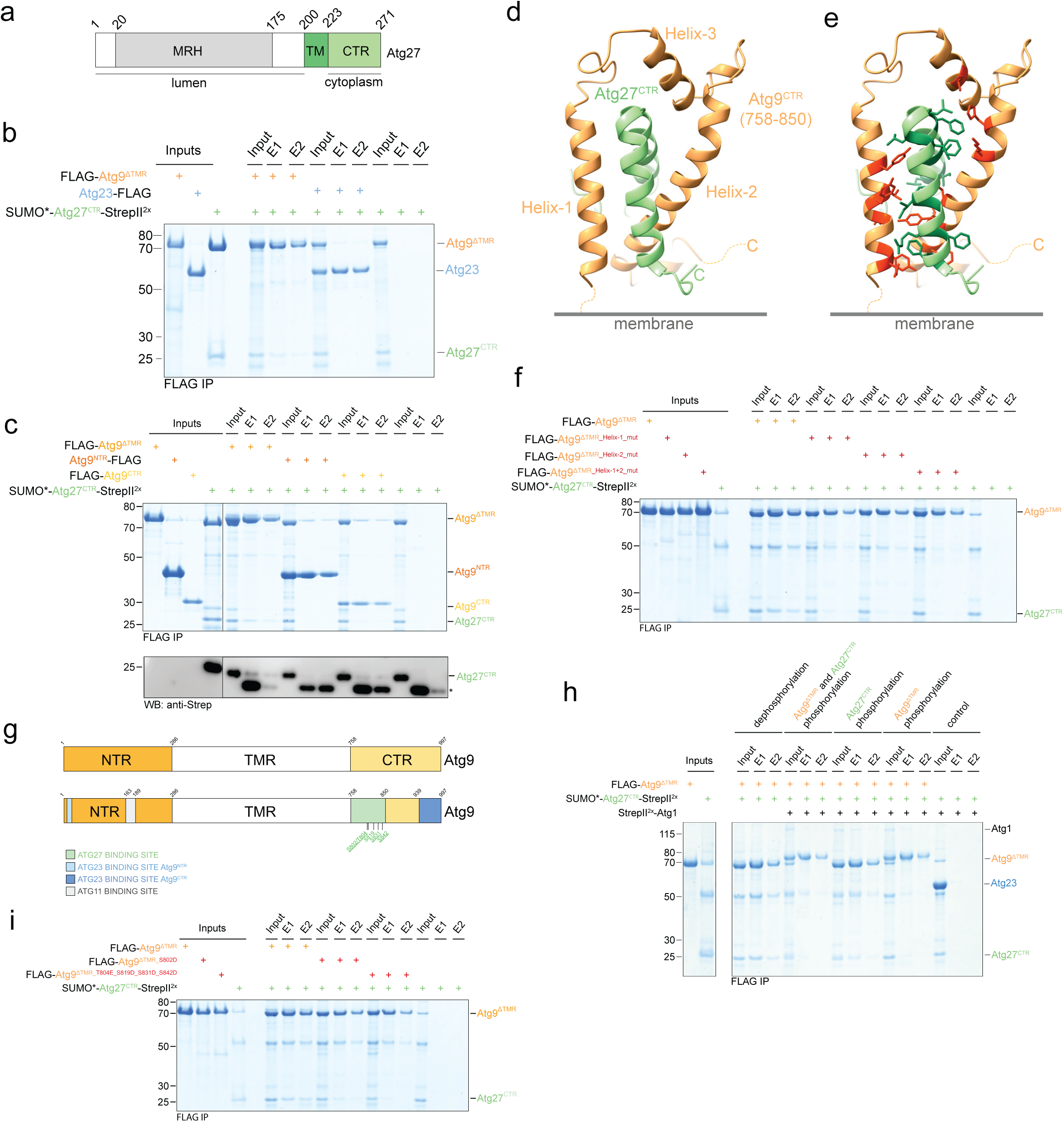
The Atg27^CTR^ interacts with the Atg9^CTR^, and this interaction is inhibited by Atg1-mediated phosphorylation of the Atg9^CTR^. **a**, Schematic domain overview of *S. cerevisiae* Atg27. Atg27 contains a luminal N-terminal mannose 6-phosphate receptor homology (MRH) domain, a single transmembrane helix (TM), and a C-terminal region (CTR) that faces the cytoplasm. **b**, The Atg27^CTR^ interacts with the Atg9^ΔTMR^ but not Atg23. FLAG-tagged Atg9^ΔTMR^ or Atg23 were mixed with SUMO*-Atg27^CTR^-StrepII^2x^ and immobilized using FLAG resin. After washing, bound proteins were eluted, and input and elution (E) samples analyzed by SDS-PAGE and Coomassie staining. **c**, The Atg27^CTR^ interacts with the Atg9^CTR^. FLAG-tagged Atg9 constructs (Atg9^NTR^, Atg9^CTR^ and Atg9^ΔTMR^) were incubated with SUMO*-Atg27^CTR^-StrepII^2x^ and immobilized using FLAG resin. After washing, bound proteins were eluted, and both input and elution (E) samples were analyzed by SDS-PAGE (top) and Western blotting using an anti-Strep antibody (bottom). The asterisk denotes the FLAG antibody light chain. **d**, AF3 model of the Atg27– Atg9 interaction. The interaction between the Atg27^CTR^ and Atg9^CTR^ was predicted using AF3 (only the interacting region of Atg9, amino acids 758-850, is shown). **e**, The AF3 model suggests a hydrophobic interface between the Atg9^CTR^ and Atg27^CTR^. AF3 model as shown in Fig. 6d with hydrophobic residues involved in the interaction highlighted in dark green (Atg27) and red (Atg9). **f**, The Atg27^CTR^-Atg9^CTR^ interaction is largely mediated by hydrophobic interactions. FLAG-tagged Atg9^CTR^ variants with alanine substitutions at critical interface residues within helix 1, helix 2, or both (Atg9^ΔTMR_Helix-1_mut^: _Atg9ΔTMR_Y762A_V763A_L766A_V769A_Y772A, Atg9ΔTMR_Helix-2_mut: Atg9ΔTMR_H818A_K821A_L829A_Y832A_V833A_F835A and_ Atg9^ΔTMR_Helix-1+2_mut^) were incubated with SUMO*-Atg27^CTR^-StrepII^2x^ and FLAG resin. After washing, bound proteins were eluted, and both input and elution (E) samples were analyzed by SDS-PAGE and Coomassie staining. **g**, Schematic domain overview of *S. cerevisiae* Atg9. The diagram highlights the newly identified Atg27 and Atg23 binding sites, along with the previously reported Atg11 binding site^31^. Previously reported phosphorylation sites within the Atg27 binding region are highlighted in green, with Atg1-dependent sites identified *in vivo* shown underlined. **h**, Atg1-dependent phosphorylation of the Atg9^ΔTMR^ inhibits Atg27^CTR^ binding. FLAG-tagged Atg9^ΔTMR^ and/or SUMO*-Atg27^CTR^-StrepII^2x^ were phosphorylated using substoichiometric amounts of recombinant Atg1. ATP was depleted using apyrase prior to combining the indicated phosphorylated and non-phosphorylated proteins. As a control, the Atg9^ΔTMR^ and the Atg27^CTR^ were dephosphorylated using -phosphatase (-PP). The Atg9^ΔTMR^ was immobilized using FLAG resin, and after washing, bound proteins were eluted, and input and elution (E) samples analyzed by SDS-PAGE and Coomassie staining. **i**, Phosphomimicking mutations in the Atg27^CTR^ binding region in Atg9 significantly reduce Atg27^CTR^ binding. FLAG-tagged Atg9^ΔTMR^ and two phosphomimicking Atg9^ΔTMR^ mutants (Atg9^ΔTMR_S802D^ and Atg9^ΔTMR_T804E_S819D_S831D_S842D^) were incubated with SUMO*-Atg27^CTR^-StrepII^2x^. Proteins were immobilized using FLAG resin. After washing, bound proteins were eluted, and input and elution (E) samples analyzed by SDS-PAGE and Coomassie staining.

Consistent with previous yeast-two-hybrid experiments^28^, the Atg27^CTR^ did not directly interact with Atg23 in the absence of Atg9 (**Fig. 6b**). However, the Atg27^CTR^ showed a direct interaction with the Atg9^ΔTMR^ (**Fig. 6b**). Further analysis revealed that binding to the Atg27^CTR^ is mediated by the Atg9^CTR^, rather than the Atg9^NTR^ (**Fig. 6c**). Using AF3, we obtained a model with reasonably high confidence for the observed Atg9^CTR^-Atg27^CTR^ interaction, suggesting that the Atg27^CTR^ binds to the region immediately C-terminal to the Atg9 transmembrane domain, encompassing residues 758 to 850 (**Fig. 6d** and **Extended Data Fig. 6b-c**). In this model, both the Atg27^CTR^ and Atg9^758–850^ adopt helix–loop–helix motifs that interlock with one another through stabilizing hydrophobic interactions. Notably, the Atg27 binding region in the Atg9^CTR^ additionally features a short third helix (helix 3) that connects the two main helices, helix 1 and helix 2 (**Fig. 6d-e**). Helix 3 also contains serine 802 (S802) which is predicted to form a hydrogen bond with glutamate 252 (E252) in Atg27 (**Extended Data Fig. 6d**).

To probe this model, we generated Atg9^ΔTMR^ constructs in which key hydrophobic residues in each of the two Atg9 *a*-helices predicted to mediate the interaction with the Atg27^CTR^ were mutated to alanine (Atg9^ΔTMR_Helix-1_mut^ and Atg9^ΔTMR_Helix-2_mut^). Consistent with the AF3 model, both Atg9^ΔTMR^ helix mutants had a significantly reduced affinity for the Atg27^CTR^ compared to wild-type Atg9^ΔTMR^ (**Fig. 6f**). Moreover, a construct in which both helices were mutated to alanine (Atg9^ΔTMR_Helix-1+2_mut^) almost fully lost its ability to bind the Atg27^CTR^ (**Fig. 6f**), further supporting the AF3 model and the Atg27-interacting region within the Atg9^CTR^ (**Fig. 6g**). Taken together, these findings indicate that the Atg23- and Atg27-interacting regions in the Atg9^CTR^ are spatially distinct (**Fig. 6g**), suggesting that the Atg9^CTR^ can simultaneously bind Atg27 and Atg23.

### Atg1-mediated phosphorylation of the Atg9^CTR^ leads to the dissociation of the Atg27^CTR^

Finally, to investigate how this interaction is regulated at the site of autophagosome formation, we tested whether the Atg9-Atg27^CTR^ interaction is also influenced by Atg1-mediated phosphorylation. Indeed, simultaneous phosphorylation of both Atg9^ΔTMR^ and Atg27^CTR^ almost completely disrupted the interaction (**Fig. 6h**). To determine whether this effect was due to phosphorylation of Atg9, Atg27, or both, we individually phosphorylated Atg9^ΔTMR^ and the Atg27^CTR^ using recombinant Atg1. This revealed that Atg1-dependent phosphorylation of the Atg9^ΔTMR^, rather than the Atg27^CTR^, regulates the interaction (**Fig. 6h**). This is consistent with previous reports showing that Atg1-dependent phosphorylation sites are exclusively found on Atg9 and not Atg27 *in vivo*^22–24^.

We therefore focused on Atg9 to identify the phosphorylation sites that regulate its interaction with Atg27. Serine 802 (S802), threonine 804 (T804), serine 819 (S819), serine 831 (S831) and serine 842 (S842) had previously been shown to be phosphorylated in cells, with all except S819 confirmed as Atg1-dependent^22–24,38^ (**Fig. 6g**). Our AF3 model suggests that S802 in Atg9 forms a hydrogen bond with glutamate 252 (E252) in the Atg27^CTR^ (**Extended Data Fig. 6d**). To test the effect of phosphorylation at this site, we introduced the corresponding phosphomimicking mutation in Atg9^ΔTMR^ (Atg9^ΔTMR_S802D^) and observed a significant decrease in Atg27^CTR^ binding (**Fig. 6i**). This result further supports the AF3 model and provides additional evidence that Atg1-mediated phosphorylation destabilizes the Atg9-Atg27 interaction as phosphorylation of S802 would interfere with hydrogen bond formation thus weakening the interaction.

To assess whether phosphorylation at additional sites affects Atg27 binding, we generated another phosphomimicking mutant in which all other previously identified Atg1-dependent phosphorylation sites and S819 were mutated to either glutamate or aspartate (Atg9^ΔTMR_T804E_S819D_S831D_S842D^). The resultant Atg9 mutant almost completely lost its affinity for the Atg27^CTR^ (**Fig. 6i**), suggesting that Atg1-mediated phosphorylation of S802 and additional site(s) within the newly identified Atg27 binding site regulate the interaction between the cytoplasmic C-termini of Atg9 and Atg27.

## Discussion

Our findings propose a molecular mechanism that clarifies Atg23’s previously suggested role in Atg9-containing vesicle formation^6,29^. Based on previous molecular dynamics simulations, it was proposed that Atg9 clustering drives the formation of dome-like structures with a curvature like that of Atg9-containing vesicles^16^. However, the mechanism by which such clustering is achieved remained unclear. Our data suggest that Atg23 may serve as the missing link: through simultaneous interactions with both the N- and C-terminal regions of Atg9, Atg23 mediates efficient clustering of Atg9 in the Golgi, thereby promoting vesicle budding. Consistent with this model, an Atg9 mutant unable to interact with Atg23 failed to promote bulk autophagy (**Fig. 5c** and **Extended Data Fig. 5c**) and did not localize to the PAS (**Fig. 5d** and **Extended Fig. 5d**). This mechanism may also explain why Atg9 overexpression can bypass the requirement for Atg23^29^, as the increase in Atg9 concentration could promote Atg9 clustering independently of Atg23.

Our data indicate that one Atg23 homodimer can bind two Atg9 termini simultaneously (**Fig. 3e**). This bivalent binding mode suggests that Atg23 can crosslink different Atg9 trimers at the Golgi, facilitating clustering. The different affinities of Atg23 for the Atg9^NTR^ and Atg9^CTR^ may further influence this process. If both interactions were of equal affinity, Atg23 might preferentially bind the termini of a single Atg9 trimer, limiting its ability to mediate inter-trimer clustering. However, given the likely asymmetric binding preferences for the Atg9^NTR^ and Atg9^CTR^ and the uneven number of C-termini per trimer, at least one Atg23 dimer may be biased towards binding in trans - linking the C-terminus of one Atg9 trimer to another trimer. The N-terminal regions of Atg9 likely provide additional anchor points to further enhance three-dimensional clustering to induce membrane curvature and promote Atg9-containing vesicle budding, which is consistent with our finding that Atg23 needs to interact with both the Atg9^NTR^ and Atg9^CTR^ to promote efficient bulk autophagy (**Fig. 5a-b**). Moreover, the recently suggested membrane-binding properties of Atg23^32^ may also play a role in vesicle budding. Indeed, the Atg23 residues implicated in membrane association^32^ are positioned away from the Atg9 binding site in Atg23, allowing Atg23 to interact simultaneously with membranes and the Atg9^NTR^ and/or Atg9^CTR^, which explains Atg23’s selectivity for Atg9-positive membranes such as the Golgi.

The predicted extended and rigid structure of the Atg23 homodimer^32^ (**Fig. 2a** and **Extended Data Fig. 2a-b**), with Atg9-binding sites located on opposite ends, appears well-suited for this bridging function. Such a geometry would favour inter-trimer crosslinking over intra-trimer binding, enhancing Atg9 clustering efficiency. Binding of the Atg27^CTR^ to the Atg9^CTR^ could influence this process by reducing the hydrodynamic reach of the Atg9 C-terminus by decreasing the length of the largely disordered C-terminus from roughly 240 amino acids (amino acids 758-997) to 138 amino acids (amino acids 850 to 997). This may further bias the extended Atg23 dimer towards engaging Atg9 trimers in *trans*rather than *cis*, thereby promoting Atg9 clustering and vesicle budding.

Our findings further indicate that Atg1-mediated phosphorylation significantly weakens Atg9 binding to both Atg23 and the C-terminal region of Atg27 and may impair interaction with Atg27 more broadly. Given that Atg1 kinase activity is high at the PAS during bulk autophagy due to the Atg13- and Atg17-dependent clustering of the Atg1 complex and resultant activation of Atg1^40^, this suggests that the protein-protein interactions between Atg9, Atg23 and Atg27 are extensively remodelled upon tethering of Atg9-containing vesicles at the PAS.

We show that the phosphorylation-dependent destabilization of the Atg9-Atg23 interaction is physiologically significant (**Fig. 5e-f**). It is tempting to speculate that Atg23 binding to Atg9 may sterically interfere with the potential fusion of Atg9-containing vesicles at the PAS. Moreover, Atg9 clustering is thought to induce strong membrane curvature^16^, which may be at least in part incompatible with autophagosomal membrane formation. Thus, release of Atg23 from Atg9 could be a prerequisite for phagophore formation.

Atg1-mediated phosphorylation might furthermore prevent the rebinding of Atg23 and Atg27 to Atg9 along the growing autophagosomal membrane, where Atg1 kinase activity is thought to be particularly high - likely due to limited co-localization with protein phosphatases^41^. This may be particularly important to avoid undesired Atg23-induced membrane deformations during phagophore expansion^42^. Moreover, the timely release of Atg23 and the Atg27^CTR^ may be required to enable other protein-protein interactions in a highly regulated spatiotemporal manner. Disruption of the Atg9-Atg23 and Atg9-Atg27 interactions could expose otherwise occluded sites on Atg9 and the Atg27^CTR^ specifically at the PAS. These sites could then recruit downstream autophagy effectors, such as the Vps34^Atg14^ complex^43^, in a likely phosphorylation-dependent manner. Atg23 and Atg27^CTR^ binding to Atg9 may also inhibit the assembly of the Atg9-Atg2-Atg18 complex on the leading edge of the phagophore^44,45^, potentially interfering with downstream steps in autophagosome biogenesis such as lipid shuttling. Finally, persistent association of Atg23 and Atg27 with Atg9 along the expanding autophagosomal membrane could sterically prevent phagophore access by other autophagy-related proteins (e.g. Atg8-interacting motif containing proteins).

Finally, the phosphoregulation of the Atg9-Atg23 interaction may also serve a spatial regulatory role, restricting the formation and budding of Atg9-containing vesicles to the Golgi, where Atg1 is not thought to localize. By coupling the dissociation of Atg23 from Atg9 to sites of high Atg1 kinase activity, such as the PAS, cells ensure that the remodelling of Atg9 interactions occurs in a tightly controlled, location-specific manner.

Taken together, our work lays the foundation for a deeper understanding of the complex mechanisms governing autophagosome formation. It provides a comprehensive protein–protein interaction and regulatory framework underlying autophagy initiation that will inform and guide future investigations into the mechanism of autophagosome formation.

## Acknowledgments

We would like to thank the Francis Crick Institute Structural Biology, Advanced Light Microscopy and Proteomics Scientific Technology Platforms (STPs) and the Schreiber lab for scientific discussions. The Schreiber lab is supported by The Francis Crick Institute, which receives its core funding from Cancer Research UK (CC2064), the UK Medical Research Council (CC2064), and the Wellcome Trust (CC2064), as well as by a UKRI (EPSRC, BBSRC and MRC) Prosperity Partnership award in collaboration with AstraZeneca (EP/X025357/1).

## Author Contributions

The study was conceptualized by A.S. with input from E.K. and C.O.S. Cloning was done by C.D., C.O.S., E.K., A.S. and X.M. Protein expression and purification was done by C.D., E.K., and C.O.S. Yeast strains were generated by C.O.S., X.M., C.D., A.S. and E.K. The biochemical analysis was carried out by E.K. and C.O.S. The MS analysis was carried out by S.M., T.A. and M.S. and the ITC experiments by E.K. and S.K. Pho8Δ60 assays were carried out by E.K. and X.M. with help from C.D. and mass photometry experiments by A.O. Fluorescence microscopy experiments were carried out by R.D., E.K. and C.O.S. The original manuscript draft was assembled by E.K and A.S with input by C.O.S. Figures were prepared by E.K, C.O.S., A.O, S.M. and A.S. Reviewing and editing of the manuscript was done by E.K., C.O.S. and A.S. Funding was acquired by A.S.

## Materials and Methods

Saccharomyces cerevisiae (S. cerevisiae) strains and media

All yeast strains used in this study are derived from *S. cerevisiae* strain BY4741 (MATa, his3Δ1, leu2Δ0, met15Δ0, ura3Δ0) and are summarized in **Supplementary Table 1**. Yeast cells were grown in YPD (1% yeast extract, 2% peptone and 2% glucose) or synthetic complete (SC) medium (0.67% yeast nitrogen base with amino acids and 0.5% glucose) at 150 rpm and 30°C. Starvation experiments were carried out by growing yeast in nitrogen starvation (SD-N) medium (0.17% yeast nitrogen base without amino acids and ammonium sulphate and 2% glucose).

### *Escherichia coli* (*E. coli*) strains and media

*E. coli* strains (DH5, BL21-CodonPlus (DE3)-RIL and DH10Multibac) were grown in Terrific Broth (TB) medium at 180 rpm and 37°C.

### Insect cells and media

Insect cells (Sf9 and High Five cells; Thermo Fisher Scientific) were grown in Sf-900 II SFM medium (GIBCO) supplemented with 0.1X Penicillin-Streptomycin-Glutamine (GIBCO) at 140 rpm and 27°C.

### Cloning

All *S. cerevisiae* genes were PCR amplified from genomic DNA. PCR products were cloned into the pCR-Blunt II-TOPO plasmid (Thermo Fisher Scientific) and subcloned into the plasmids listed in **Supplementary Table 2** using restriction enzyme digestion and ligation. Restriction sites, mutations and tags were introduced by In-Fusion cloning (TaKaRa Bio) or PCR. PCR products were gel extracted using the GeneJET PCR Purification Kit (Thermo Fisher Scientific). Plasmids were transformed into *E. coli* DH5¤, isolated using the GeneJET Plasmid Miniprep Kit (Thermo Fisher Scientific) and sequence verified.

### Plasmids used for *S. cerevisiae* strain construction

All plasmids used for *S. cerevisiae* strain generation are pCR-Blunt II-TOPO derivatives and are listed in **Supplementary Table 2**. They contain the gene-specific promoter (∼ 500 bp upstream of the gene-specific start codon) as a region of homology, the mutated open reading frame or the gene fusion construct, the terminator sequence (∼150-300 bp downstream of the gene-specific stop codon), the selection cassette and a second region of homology downstream of the terminator sequence (∼300-500 bp). Gene deletions were generated by fully replacing the target gene with the indicated selection cassettes. The plasmids were linearized and transformed into the yeast strains of interest and the final yeast strains were verified by PCRs and Sanger sequencing.

### Plasmids used for bacterial protein expression

All genes were cloned into pET17b or pET17b-SH-SUMO* plasmids (SH: His_6_-StrepII^2x^). The resultant plasmids are listed in **Supplementary Table 2**.

### Baculovirus generation

Genes were cloned into pFBDM transfer plasmids. The resultant plasmids (listed in **Supplementary Table 2**) were transformed into DH10Multibac cells and bacmids were isolated using isopropanol precipitation. GeneJuice Transfection Reagent (Sigma) was used to transfect Sf9 cells with bacmids. Viruses were amplified up to the P3 stage in Sf9 cells using a multiplicity of infection (MOI) of 0.1, according to standard protocols.

### Protein expression in bacteria

For bacterial protein expression, plasmids were transformed into BL21-CodonPlus (DE3)-RIL cells (Agilent). Cells were grown at 37°C in TB medium supplemented with ampicillin (100 mg/mL) and chloramphenicol (25 mg/mL) until they reached an OD_600_ of 0.8-1.0. Cells were moved on ice and protein expression was induced with 0.5 mM isopropyl-b-D-1-thiogalactopyranoside (IPTG). Protein expression was carried out overnight at 18°C. Cells were harvested at 4000 rpm (Beckman Coulter J6-MC Centrifuge) for 10 min at 4°C.

### Protein expression in insect cells

High Five cells were grown to a density of ∼2 × 10^6^ cells/mL and infected with a P3 baculovirus using a MOI greater than 2. Protein expression was carried out at 27°C for 60-72h. Cells were harvested at 3000 rpm (Beckman Coulter J6-MC Centrifuge) for 10 min at 4°C.

### Purification of Autophagy-related (Atg) proteins and protein complexes

*S. cerevisiae* Atg proteins were purified at 4°C. Bacterial or insect cell pellets were resuspended in pre-cooled Lysis buffer containing 50 mM Tris-HCl pH 8.5, 5% glycerol, 300 or 200 mM NaCl (for monomeric proteins or protein complexes, respectively), 2 mM DTT, 4 mM EDTA, 0.2 mM PMSF, 10 mM leupeptin, 10 mM pepstatin A, 1 mM benzamidine, cOmplete EDTA-free Protease Inhibitor Cocktail (1 tablet per 50 mL, Roche) and Pierce Universal Nuclease (6 µL per 100 mL, Thermo Fisher Scientific). The Lysis buffer used for bacterial protein purifications was supplemented with lysozyme (100 mg/mL). Cells were lysed by sonication and spun at 20,000 rpm for 1 h using a Beckman Coulter JA-20 rotor.

### Affinity purifications

Supernatants were loaded onto a 5 mL StrepTactin column (QIAGEN) or 1 mL StrepTactin column (IBA Lifesciences) pre-equilibrated with Wash buffer composed of 50 mM Tris-HCl pH 8.0, 300 or 200 mM NaCl (for monomeric proteins or protein complexes, respectively), 5% glycerol and 2 mM DTT. The column was washed with 10-20 column volumes (CV) Wash buffer before proteins were eluted with 5 CV Wash buffer containing 2.5 mM desthiobiotin.

Depending on the experiment, tags were removed from eluted proteins by overnight incubation at 4 °C with PreScission (3C) protease at a 1:50 protease-to-protein molar ratio, following dilution to a final salt concentration of approximately 100–200 mM NaCl. Proteins were subsequently purified by ion exchange chromatography or concentrated using Amicon Ultra concentrators (Sigma) and subjected to size exclusion chromatography (SEC).

### Ion exchange chromatography

Elution samples from the affinity purification were diluted to a final salt concentration of ∼100-150 mM NaCl and subjected to ion exchange chromatography. Proteins were purified by anion exchange chromatography using a ResQ column (GE Healthcare), applying a salt gradient from 50 to 700 mM NaCl (basic ResQ buffer composition: 20 mM HEPES-NaOH pH 8.0, 5% glycerol and 2 mM DTT). Protein-containing fractions were pooled, and if 3C cleavage had been performed, the sample was passed back over a StrepTactin column to remove the tag and any uncleaved protein. This step was omitted if further purification by SEC was required. The proteins were then concentrated and either snap-frozen or subjected to additional purification by SEC.

### Size exclusion chromatography (SEC)

Samples were loaded on a SEC column pre-equilibrated in SEC buffer (20 mM HEPES-NaOH pH 7.4, 180 mM NaCl, 5% glycerol and 2 mM DTT). If 3C cleavage had been performed, a StrepTactin pass-back column was connected at the end of the SEC column to remove the cleaved tag and any uncleaved protein. Protein-containing fractions were pooled, concentrated and snap-frozen.

### Pulldown assays

Strep, Myc and FLAG pulldown experiments were carried out using either StrepTactin Superflow Plus (QIAGEN), Anti-c-Myc Agarose (Thermo Fisher Scientific) or Anti-FLAG M2 affinity resin (Sigma). Prior to sample addition, the resin was sequentially washed with each 10 bed volumes (BV) of 1X TBS (0.05 M Tris-HCl pH 7.6 and 0.15 M NaCl), 0.1 M glycine-HCl pH 3.4 and twice 1X TBS before the resin was equilibrated in Pulldown buffer (20 mM HEPES-NaOH pH 7.5, 150 mM NaCl). Bait and prey proteins were typically mixed in a 1:2 molar ratio and incubated with the resin for 30 min at room temperature (RT). Beads were washed three times with Pulldown buffer and bound proteins eluted using Pulldown buffer supplemented with either 2.5 mM desthiobiotin, 300 µg/mL Myc peptide (GenScript) or 300 µg/mL 3xFLAG peptide (Sigma). Pulldowns were analysed by Western blotting or SDS-PAGE using either Quick Coomassie Stain (Neo Biotech) or Sypro Ruby Protein Gel Stain (Thermo Fisher Scientific).

To probe interactions of non-phosphorylated proteins, proteins were mixed in Pulldown buffer in the presence of Lambda Protein Phosphatase (NEB) and incubated for 30-45 min at RT before the pulldown experiment. To assess the effect of Atg1-mediated phosphorylation, the target proteins were incubated in Pulldown buffer supplemented with 5 mM MgCl,, 1 mM ATP, and 1× PhosSTOP protein phosphatase inhibitors (Roche) to prevent dephosphorylation by any co-purified phosphatases. Recombinant Atg1 kinase was added at a 1:50 molar ratio of kinase to substrate. Protein phosphorylation was allowed to proceed for 45 min at 30°C prior to the pulldown experiments. If only a subset of the pulldown components required phosphorylation, apyrase was added after completion of the phosphorylation reaction, and the mixture was incubated for an additional hour at 30 °C to deplete residual ATP before introducing the remaining protein(s).

### Western blotting (WB)

Proteins were separated by SDS-PAGE and transferred to Immobilon-FL PVDF membranes (Sigma) using a Criterion Blotter (BioRad). Membranes were incubated either with monoclonal anti-Strep (BioRad), anti-c-Myc (Cell Signalling Technology), anti-FLAG M2 (Sigma) or anti-PGK1 (22C5D8, Invitrogen) primary antibodies. After washing, membranes were incubated with an anti-mouse IgG secondary antibody (BioRad). Immobilon Classico Western HRP Substrate (Sigma) or Clarity Western ECL Substrate (BioRad) was used to visualise the HRP-conjugated secondary antibody. Detection of proteins was performed using an Amersham ImageQuant 800 biomolecular imager (Cytiva).

### Isothermal titration calorimetry (ITC)

Atg23, Atg9^NTR^-FLAG and FLAG-Atg9^CTR^ were dialysed against the ITC buffer (20 mM HEPES-NaOH pH 7.5, 150 mM NaCl, 1 mM TCEP). Final concentrations were determined by UV absorbance measurements with a V-760 UV-Visible Spectrophotometer (Jasco), using the extinction coefficients s_280_ = 25440 M^-1^ cm^-1^ (Atg23), s_280_ = 5960 M^-1^ cm^-1^ (Atg9^NTR^-FLAG) and s_280_ = 33350 M^-1^ cm^-1^ (FLAG-Atg9^CTR^). Binding parameters were determined using a MicroCal PEAQ-ITC instrument (Malvern Panalytical). To probe the Atg9^NTR^-Atg23 interaction, Atg9^NTR^-FLAG was loaded into the sample cell at 45 µM and Atg23 into the syringe at a monomer concentration of 405 µM (repeats 1 and 2). To probe the Atg9^CTR^-Atg23 interaction, FLAG-Atg9^CTR^ was loaded into the sample cell at 10.0 µM (repeat 1) or 12.9 µM (repeat 2) and Atg23 into the syringe at a monomer concentration of 162 µM (repeat 1) or 184 µM (repeat 2). Atg23 was injected into the sample cell using 19 injections of 2 µL, with a reference power of 5 µcal/s and at a temperature of 25°C. Data was analysed using the One Set of Sites model in the MicroCal PEAQ-ITC Analysis Software (v1.41, Malvern Panalytical).

### Mass photometry (MP)

MP data were acquired on a Two^MP^ mass photometer instrument (Refeyn)^46^. Prior to measurements, a drop of objective immersion oil (ImmersolTM 518F Immersion Oil, ZeissTM) was added onto the objective, and a cleaned glass coverslip (No. 1.5, 24 x 50, VWR) with a gasket (CultureWell gaskets, well capacity 3-10 µL, Grace Bio-Labs) was placed on top of the objective. For automated focus adjustment, 18 µL of 0.2 µm-filtered MP buffer (20 mM HEPES-NaOH, 180 mM NaCl, 0.5 mM TCEP, 5% glycerol, pH 7.4) was added to a well. Once in focus, 2 µL of the Atg23-Myc sample diluted in the MP buffer was mixed into the well by pipetting, resulting in a 25, 50 and 75 nM measurement concentration. Particle landing data was acquired for 60 seconds per sample (2 recordings each) and analysed using DiscoverMP software (version 2024 R1, Refeyn) with in-built histogram fitting.

### Fluorescence microscopy

Fluorescence microscopy experiments were carried out as previously described^41^. Briefly, yeast strains were exponentially grown in YPD medium or starved for 4h in SD-N medium. For imaging, cells grown in YPD were pelleted, and resuspended in Synthetic Complete (SC) medium, while those starved in SD-N were kept in the same medium. Cells were diluted to an OD_600_ of 0.05 and added to P96-1.5H-N plates (Cellvis) prior to imaging. Images were acquired in a temperature-controlled environment (30°C) on an inverted wide-field Nikon Eclipse Ti2 microscope, equipped with a 100X/1.4 NA lens and a Prime 95B sCMOS camera (Teledyne Photometrics), controlled through Micro-Manager 2.0 software^47^. Z-stacks were acquired using selective band-pass filters for EGFP, BFP, and transmitted light over a range of 10 µm, with a step size of 0.25 µm. Deconvolution of the fluorescence channels was obtained by processing the datasets with scikit-image 0.18.1 python package^48^, adopting the Richardson-Lucy algorithm, with 15 iterations. Figures were prepared using Fiji^49^.

### Pho8Δ60 assays

‘‘YPD’’ samples were prepared by growing yeast cells to an OD_600_ of 0.9-1.2 in YPD medium at 30°C. ‘‘SD-N’’ samples were prepared by replacing the YPD medium with SD-N medium. Cells were washed with SD-N medium before growing them for the indicated time points in SD-N medium. Cells were harvested at 4000 rpm at RT. Cell pellets were first washed with ice-cold water and subsequently with resuspension solution (0.85% NaCl and 1 mM PMSF). Cells were processed as described previously^39^ and alkaline phosphatase activity was measured using an end-point spectrophotometric assay monitoring hydrolysis of p-nitrophenolphosphate (pNPP) to p-nitrophenol (pNP). The average and standard deviations were calculated based on at least three biological replicates.

### Hydrogen-deuterium exchange (HDX) mass spectrometry

Individual proteins (SH-SUMO*-Atg9^NTR^-FLAG, Myc-Atg9^CTR^-StrepII^2x^ and Atg23-Myc) and protein complexes (SH-SUMO*-Atg9^NTR^-FLAG/Atg23-Myc and Myc-Atg9^CTR^-StrepII^2x^/Atg23-Myc) were diluted to a final concentration of 10 μM and 5 μL of the different samples were incubated with 40 μl of D_2_O buffer at RT for 3, 30, 300 and 3000 seconds in triplicate. The labelling reaction was quenched by adding chilled 2.4% v/v formic acid in 2 M guanidinium hydrochloride and immediately frozen in liquid nitrogen. Samples were stored at -80°C prior to analysis.

The quenched protein samples were rapidly thawed and subjected to proteolytic cleavage by pepsin followed by reverse phase HPLC separation. Briefly, the proteins were passed through an Enzymate BEH immobilized pepsin column, 2.1 x 30 mm, 5 μm (Waters) at 200 μL/min for 2 min and the peptic peptides trapped and desalted on a 2.1 x 5 mm C18 trap column (Acquity BEH C18 Van-guard pre-column, 1.7 μm, Waters). Trapped peptides were subsequently eluted over 12 min using a 5-36% gradient of acetonitrile in 0.1% v/v formic acid at 40 μL/min. Peptides were separated on a reverse phase column (Acquity UPLC BEH C18 column 1.7 μm, 100 mm x 1 mm, Waters). Peptides were detected on a Cyclic mass spectrometer (Waters) acquiring over a m/z of 300 to 2000, with the standard electrospray ionization (ESI) source and lock mass calibration using [Glu1]-fibrino peptide B (50 fmol/μL). The mass spectrometer was operated at a source temperature of 80°C and a spray voltage of 3.0 kV. Spectra were collected in positive ion mode.

Peptide identification was performed by MSe^50^ using an identical gradient of increasing acetonitrile in 0.1% v/v formic acid over 12 min. The resulting MSe data were analyzed using Protein Lynx Global Server software (Waters) with an MS tolerance of 5 ppm.

Mass analysis of the peptide centroids was performed using DynamX sotware (Waters). Only peptides with a score >6.4 were considered. The first round of analysis and identification was performed automatically by the DynamX software, however, all peptides (deuterated and non-deuterated) were manually verified at every time point for the correct charge state, presence of overlapping peptides, and correct retention time. Deuterium incorporation was not corrected for back-exchange and represents relative, rather than absolute changes in deuterium levels. Changes in H/D amide exchange in any peptide may be due to a single amide or a number of amides within that peptide. All time points in this study were prepared at the same time and individual time points were acquired on the mass spectrometer on the same day.

### Structure Predictions

Structural predictions were performed using the AlphaFold 3 web server^33^. Input sequences used for the predictions are indicated in the text and corresponding figure legends. Model confidence was assessed using the provided pIDDT scores.

### Quantification and statistical analysis

As indicated in the figure legends, data are represented as mean ± standard deviation. Comparisons of two constructs in a pulldown assay were done using an unpaired, two-tailed Student’s t-test, as specified in the respective figure legends. To compare the autophagy phenotype of multiple strains in Pho8Δ60 assays, the one-way ANOVA in combination with Tukey’s multiple comparisons post-hoc test was used, as described in the figure legends.

---

**Extended Data Fig. 1:**
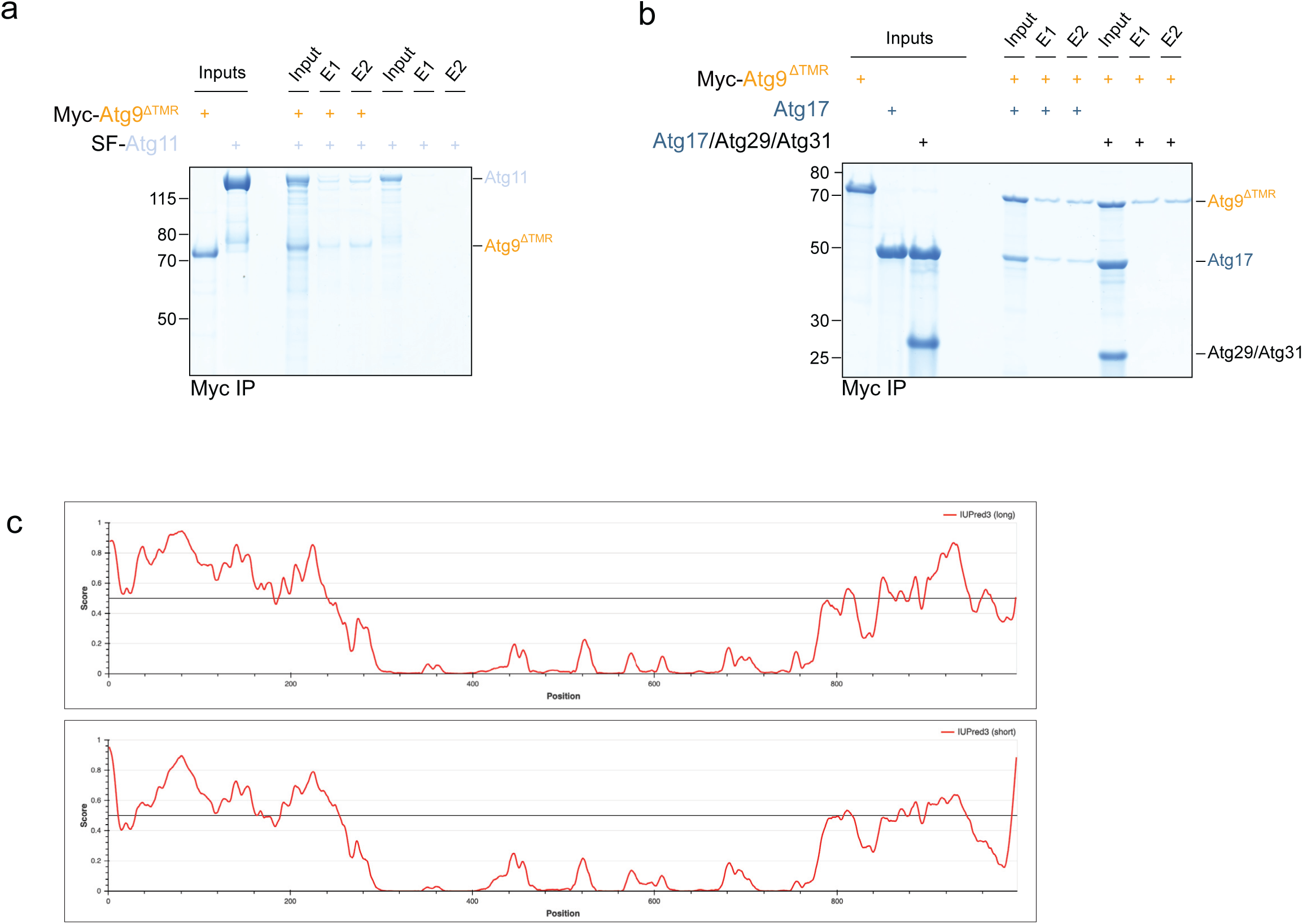
Atg23 interacts with both the N- and C-terminal regions of Atg9. **a**, Atg9^ΔTMR^ interacts with the selective autophagy-specific Atg1 complex subunit Atg11. Myc-tagged Atg9^ΔTMR^ was immobilized using Myc resin and incubated with FLAG-StrepII^2x^-tagged Atg11 (SF-Atg11). After washing the resin, bound proteins were eluted and input and elution (E) fractions were analysed by SDS-PAGE and Coomassie staining. **b**, The Atg9^ΔTMR^ interacts with Atg17, a subunit of the bulk autophagy-specific Atg1 complex; this interaction is blocked in the presence of the Atg29-Atg31 complex. Myc-tagged Atg9^ΔTMR^ was immobilized using Myc resin and incubated with Atg17 or the Atg17-Atg29-Atg31 complex. Bound proteins were eluted after washing the resin, and both input and elution (E) samples were analyzed by SDS-PAGE and Coomassie staining. **c**, Intrinsic disorder prediction of *S. cerevisiae* Atg9 using IUPred3. Scores above the 0.5 threshold indicate regions predicted to be intrinsically disordered. The analysis was performed using the IUPred3 long (top) or short (bottom) disorder prediction mode.

**Extended Data Fig. 2:**
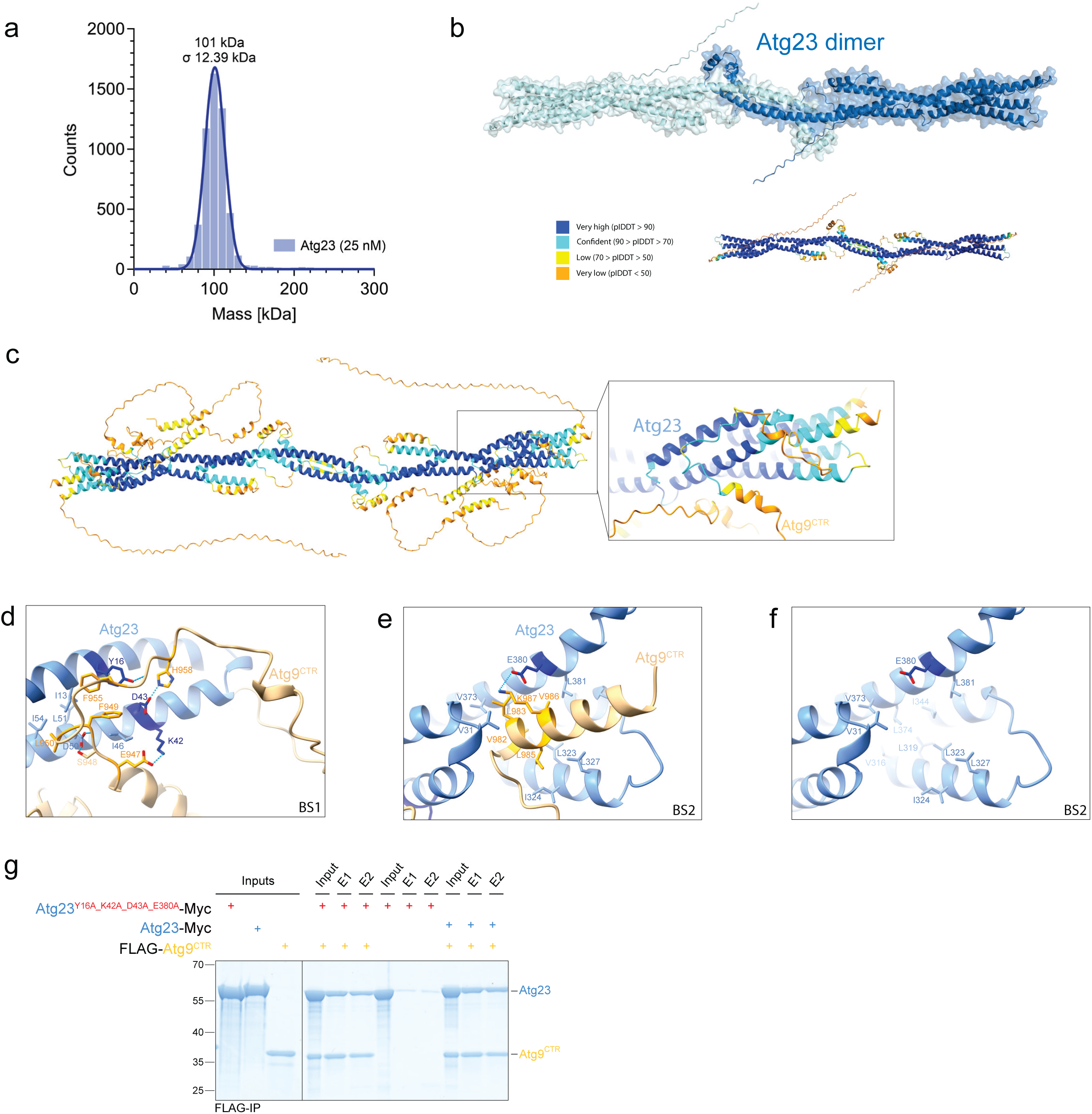
Atg23 interacts with the very C-terminus of Atg9. **a**, Mass photometry analysis of *S. cerevisiae* Atg23 demonstrates that Atg23 is a homodimer. Mass photometry histogram for Atg23-Myc. Myc-tagged Atg23 was measured at a concentration of 25 nM (n = 2). The theoretical molecular weight for a monomer and dimer is 52.7 and 105.4 kDa, respectively. **b**, AF3 model of the Atg23 homodimer. The model is color-coded by pIDDT values (bottom), with high-confidence regions (pIDDT ≥ 70) shown in dark and light blue. **c**, Prediction confidence of the AF3 Atg9^CTR^-Atg23 complex model shown in Fig. 2a. The model is color-coded according to plDDT values. The inset highlights Atg9 binding sites 1 and 2 (BS1 and BS2). **d-e**, AF3 model showing the Atg9 BS1 (panel d) and BS2 (panel e) with the residues that were mutated in the Atg23 interface mutant (Atg23^Y16A_K42A_D43A_E380A^) highlighted in dark blue. **f**, AF3 model as depicted in Extended Data Fig. 2e with only Atg23 and the critical interface residues shown. **g**, The Atg23^Y16A_K42A_D43A_E380A^ mutant still interacts with the Atg9^CTR^. The FLAG-tagged Atg9^CTR^ was immobilized using FLAG resin and incubated with Myc-tagged Atg23 or the Atg23^Y16A_K42A_D43A_E380A^ mutant. After washing the resin, bound proteins were eluted and input and elution (E) fractions were analysed by SDS-PAGE and Coomassie staining.

**Extended Data Fig. 3:**
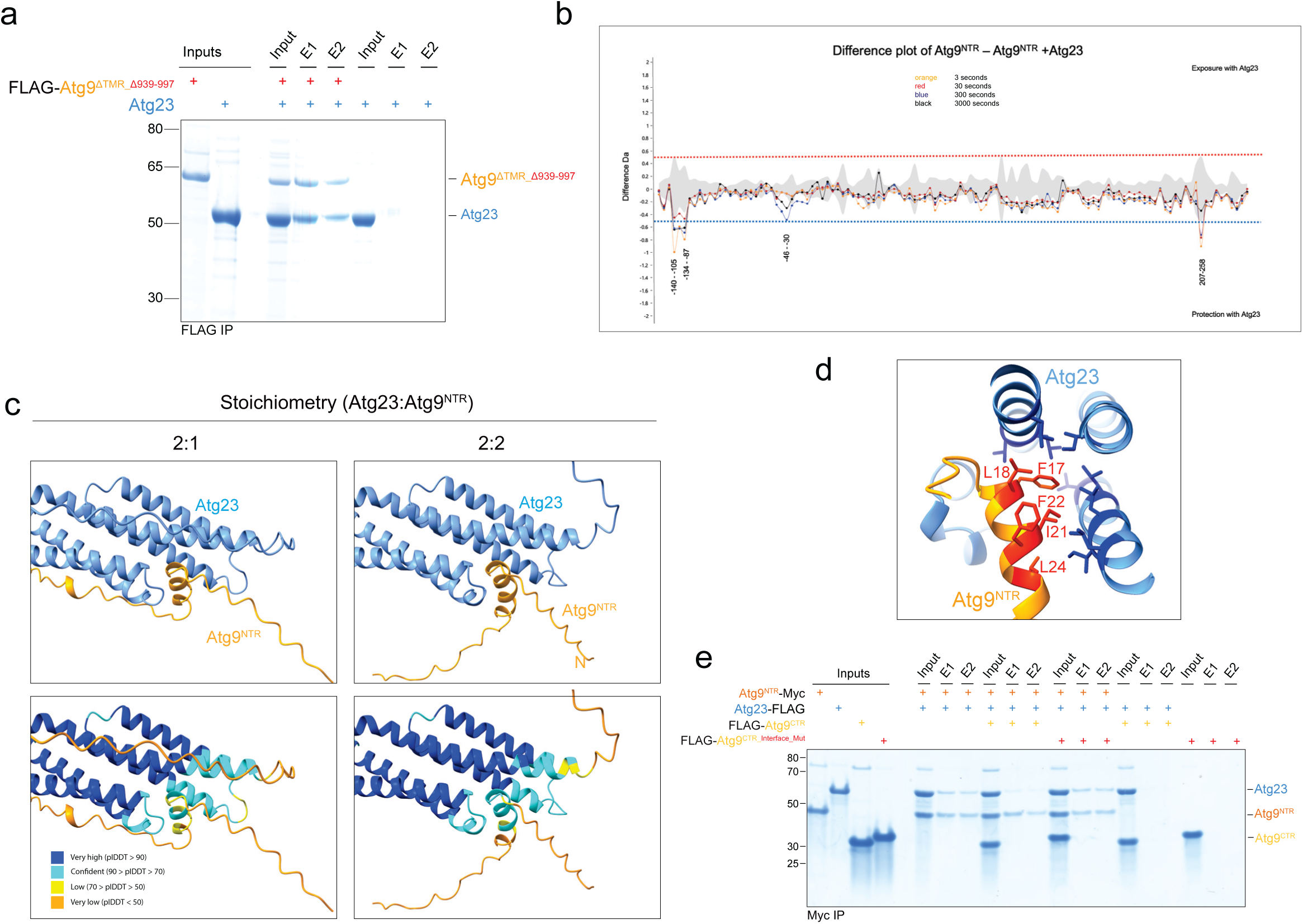
Atg23 also interacts with the Atg9^NTR^, using the same binding site it employs for binding the Atg9^CTR^. **a**, Deletion of the very C-terminus in the Atg9^ΔTMR^ does not prevent Atg23 binding. A FLAG-tagged C-terminal deletion mutant of Atg9^ΔTMR^ (Atg9^ΔTMR_Δ939-997^) was immobilized using FLAG resin and incubated with Atg23. After washing the resin, bound proteins were eluted and input and elution (E) fractions were analysed by SDS-PAGE and Coomassie staining. **b**, HDX MS difference plots comparing the Atg9^NTR^ alone and in complex with Atg23. Difference plots show hydrogen–deuterium exchange in the SH-SUMO*-tagged Atg9^NTR^ compared to that in the SH-SUMO*-Atg9^NTR^-Atg23 complex. Negative values below the significance threshold of -0.5 indicate regions that become protected upon complex formation. Deuterium uptake was measured at four time points: 3 seconds (orange), 30 seconds (red), 300 seconds (blue), and 3000 seconds (black). Negative numbers for amino acids refer to the N-terminal SH-SUMO*-tag (SH: His_6_-StrepII^2x^-tag). **c**, Zoom-in views of the predicted Atg23-Atg9^NTR^ binding interface highlighted in Fig. 3a, comparing two stoichiometries of the Atg23-Atg9^NTR^ complex: 2:1 (left panels) and 2:2 (right panels). AF3 model confidence is indicated by pIDDT color coding (lower panels). **d**, Predicted Atg9^NTR^-Atg23 interface. Hydrophobic residues in the Atg9^NTR^ (in red) and in Atg23 (in dark blue) predicted by AF3 to contribute to the Atg9^NTR^-Atg23 interface are highlighted. These residues were mutated in the Atg9^NTR_Interface_Mut^ construct (shown in Fig. 3d). **e**, The Atg9^CTR^ can outcompete the Atg9^NTR^ for binding to Atg23. Myc-tagged Atg9^NTR^ and FLAG-tagged Atg23 were incubated in the presence or absence of the FLAG-tagged wild-type Atg9^CTR^ or the interface disrupting Atg9^CTR_Interface_Mut^ mutant. The Atg9^NTR^ was immobilized using Myc resin and following washing, bound proteins were eluted. Both input and elution (E) samples were analyzed by SDS-PAGE and Coomassie staining.

**Extended Data Fig. 4:**
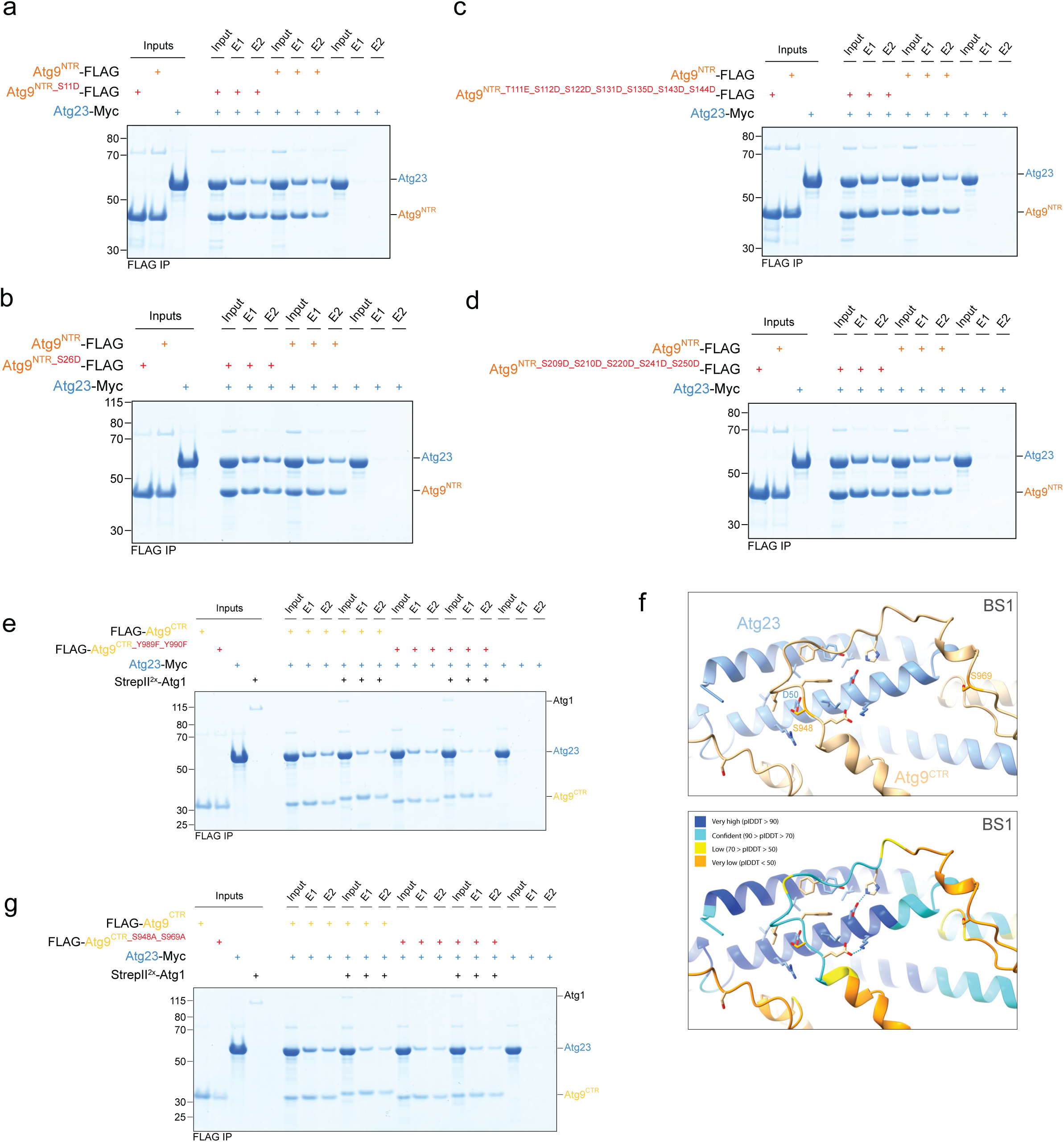
The Atg9-Atg23 interaction is inhibited by Atg1-mediated phosphorylation of Atg9. **a-d**, Phosphomimicking mutations of phosphorylation sites other than in the very N-terminus do not significantly regulate Atg23 binding. FLAG-tagged Atg9^NTR^ and the phosphomimicking mutants Atg9^NTR_S11D^ (panel **a**), Atg9^NTR_S26D^ (panel **b**), _Atg9NTR_T111E_S112D_S122D_S131D_S135D_S143D_S144D (panel_ **_c_**_), or Atg9NTR_S209D_S210D_S220D_S241D_S250D_ (panel **d**) were incubated with Myc-tagged Atg23. Proteins were immobilized on FLAG resin, washed and eluted. Input and elution (E) samples were analyzed by SDS-PAGE and Coomassie staining. **e**, Atg9 residues Y989 and Y990 are not involved in the Atg1-mediated regulation of the Atg9^CTR^-Atg23 interaction. FLAG-tagged Atg9^CTR^ or the Atg9^CTR_Y989F_Y990F^ mutant were incubated with Myc-tagged Atg23 and either phosphorylated using recombinant Atg1 or dephosphorylated using lambda phosphatase and subsequently immobilized using anti-FLAG resin. After washing, bound proteins were eluted, and both input and elution (E) samples were analyzed by SDS-PAGE followed by Coomassie staining. **f**, AF3-predicted model of Atg9 Binding Site 1 (BS1; top) and corresponding pIDDT values (bottom). S948 maps to a highly kinked segment in the Atg9^CTR^ that interacts with Atg23. Regions with pIDDT values ≥ 70 (dark and light blue) are predicted with high accuracy. **g**, Atg1-mediated phosphorylation of S948 and S969 regulates the Atg9^CTR^-Atg23 interaction. The FLAG-tagged Atg9^CTR^ or the phosphorylation-deficient Atg9^CTR_S948A_S969A^ mutant were mixed with Myc-tagged Atg23 and either phosphoryated using recombinant Atg1 or dephosphorylated using lambda phosphatase. The proteins were subsequently immobilized on anti-FLAG resin. After washing, bound proteins were eluted, and both input and elution (E) samples were analyzed by SDS-PAGE followed by Coomassie staining. Notably, the basal binding affinity of the Atg9^CTR_S948A_S969A^-Atg23 complex appears reduced, likely due to the loss of a hydrogen bond between Atg9 S948 and Atg23 D50.

**Extended Data Fig. 5:**
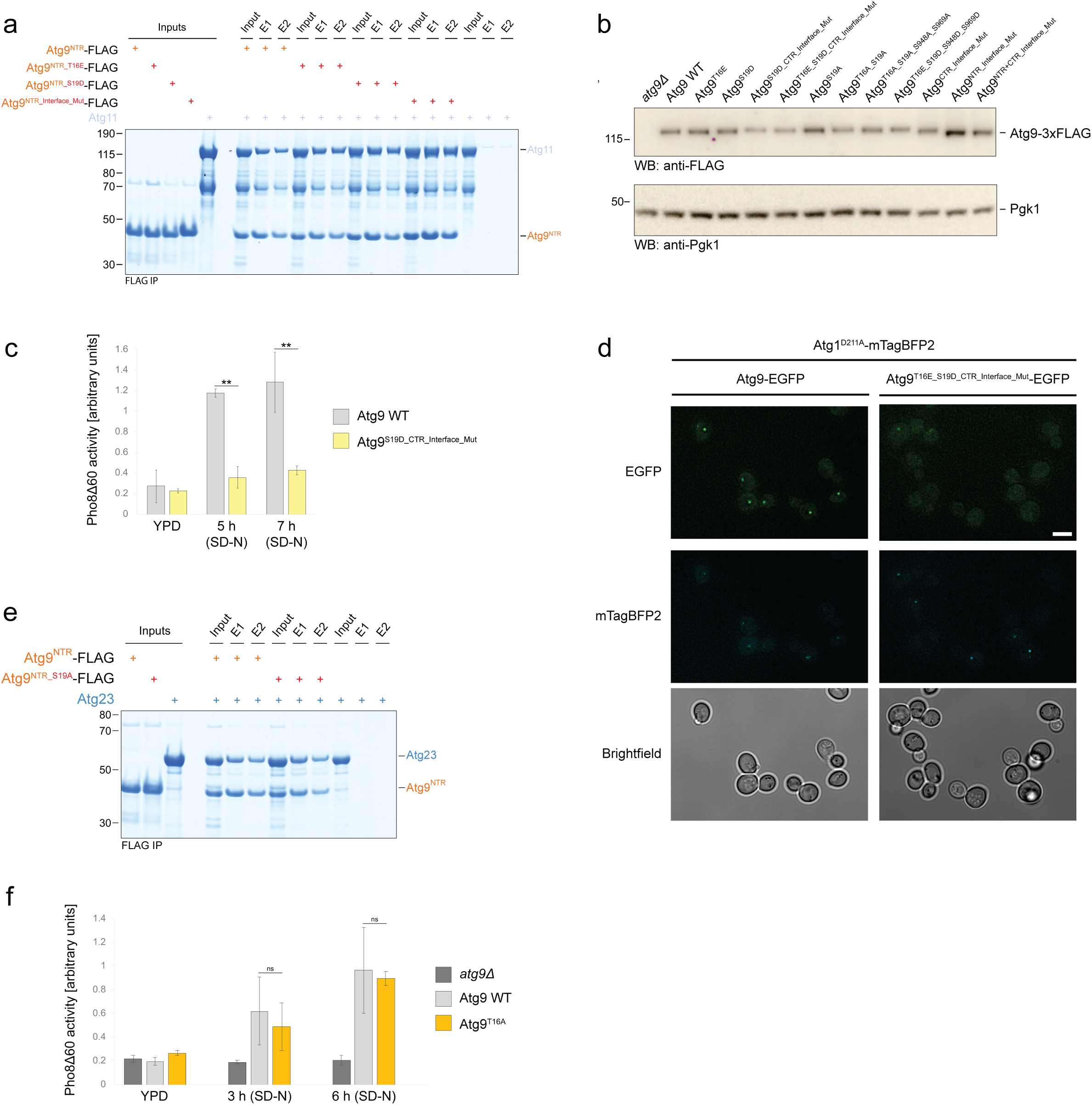
The Atg9-Atg23 interaction and its regulation by Atg1 are required for bulk autophagy. **a**, The Atg9^NTR^ phosphomimicking and interface mutants are still able to bind Atg11. The FLAG-tagged Atg9^NTR^, Atg9^NTR_T16E^, Atg9^NTR_S19D^ or Atg9^NTR_Interface_Mut^ mutants were mixed with Atg11 and immobilized using anti-FLAG resin. After washing, bound proteins were eluted, and both input and elution (E) samples were analyzed by SDS-PAGE followed by Coomassie staining. **b**, Western blot (WB) analysis of yeast strains expressing 3xFLAG-tagged wild-type Atg9 and the indicated phosphomimicking, phosphorylation-deficient, and interface mutants. **c**, The Atg9^S19D_CTR_Interface_Mut^ mutant fails to support bulk autophagy. Pho8Δ60 assay measuring bulk autophagy in *S. cerevisiae* strains expressing either wild-type Atg9 or _the Atg9S19D_CTR_Interface_Mut (Atg9S19D_E947A_F949A_L950A_F955A_H958A_V982A_L983A_L985A_V986A_K987A) mutant._ Cells were grown exponentially in YPD medium or subjected to nitrogen-starvation (SD-N) for 5 or 7 hours. Pho8Δ60-specific alkaline phosphatase activity was quantified, with mean values from three biological replicates (n = 3) presented. Error bars indicate the standard deviation. Statistical significance between yeast strains was assessed using one-way ANOVA followed by Tukey’s multiple comparisons test. **d**, The Atg9^T16E_S19D_CTR_Interface_Mut^ mutant exhibits strongly impaired PAS localization. Fluorescence microscopy analysis of cells expressing Atg1^D211A^-mTagBFP2 as a PAS marker and either EGFP-tagged wild-type Atg9 or the Atg9^T16E_S19D_CTR_Interface_Mut^ mutant. Cells were grown exponentially in nutrient-rich YPD medium and imaged in SC medium. Scale bar: 5 µm. **e**, The Atg9 S19A mutation does not affect the interaction between the Atg9^NTR^ and Atg23. The FLAG-tagged Atg9^NTR^ or the Atg9^NTR_S19A^ mutant were mixed with Atg23 and immobilized using anti-FLAG resin. After washing, bound proteins were eluted, and both input and elution (E) samples were analyzed by SDS-PAGE followed by Coomassie staining. **f**, Pho8Δ60 experiments were carried out and analysed as described in Extended Data Fig. 5c. A *S. cerevisiae* strain expressing the phosphorylation-deficient Atg9^T16A^ mutant was compared to cells expressing wild-type Atg9 and an *atg9Δ* strain. Cells were grown exponentially in YPD medium or subjected to 3 or 6 hours of nitrogen-starvation (SD-N). All Atg9 variants tested contained a C-terminal 3xFLAG tag (panels **c** and **f**).

**Extended Data Fig. 6:**
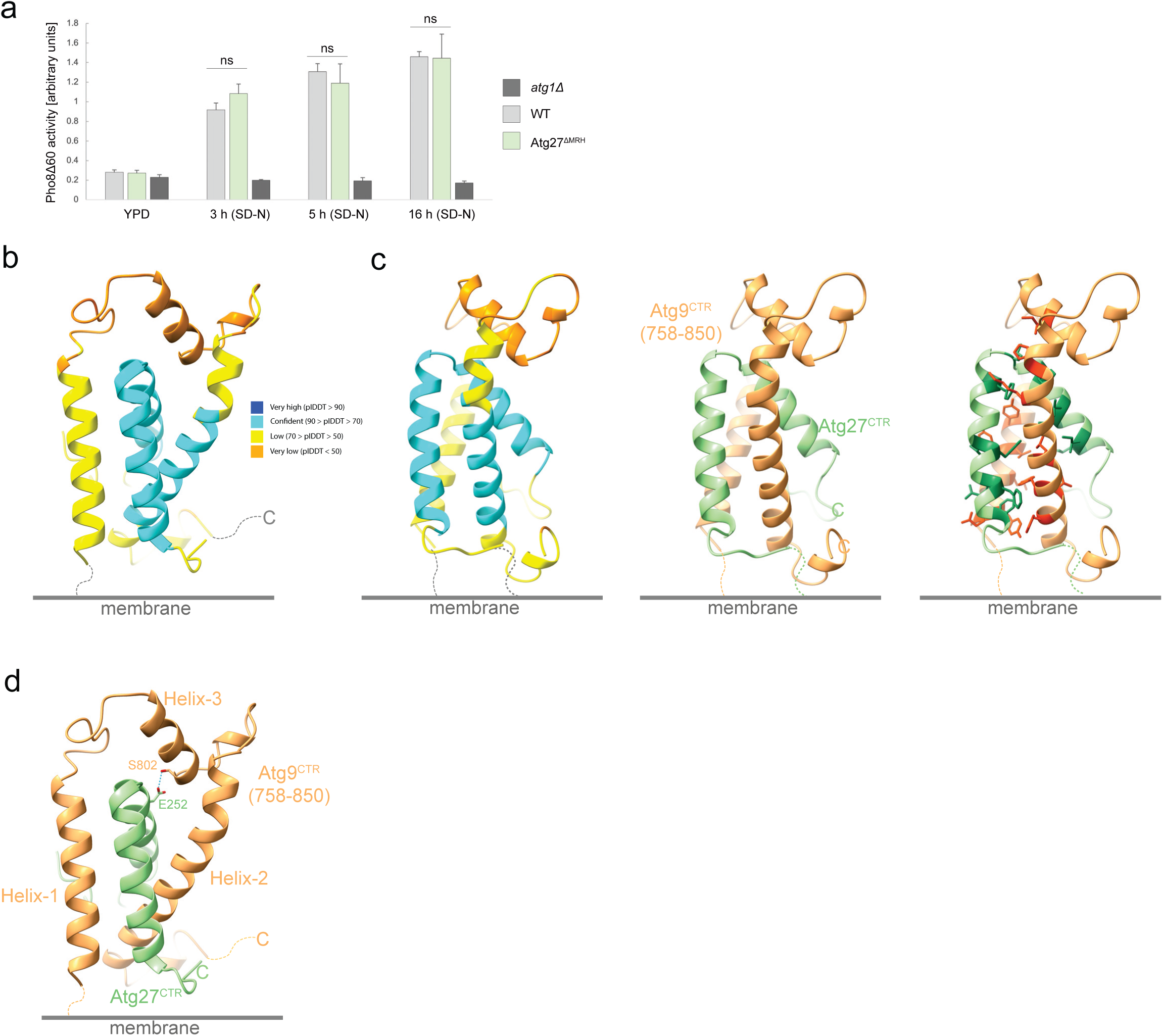
The Atg27^CTR^ interacts with the Atg9^CTR^, and this interaction is inhibited by Atg1-mediated phosphorylation of the Atg9^CTR^. **a**, The MRH domain in Atg27 is dispensable for autophagy. Pho8Δ60 assay measuring bulk autophagy in budding yeast expressing either wild-type Atg27 or an Atg27 mutant lacking the luminal MRH domain (Atg27^ΔMRH^). An *atg1Δ* strain was included as a negative control. Cells were grown exponentially in YPD medium or subjected to nitrogen-starvation (SD-N) for 3, 5 or 16 hours. Pho8Δ60-specific alkaline phosphatase activity was measured, and the mean activity (n = 3) is shown with error bars representing the standard deviation. A one-way ANOVA in combination with Tukey’s multiple comparisons test was used to assess the statistical significance of differences in Pho8Δ60 activity between strains. **b**, The AF3 model shown in Fig. 6d-e is color-coded according to the pIDDT values. The accuracy of the prediction for regions with pIDDT values ≥ 70 (dark and light blue) is high. **c**, Different view of the AF3 model shown in Fig. 6d-e. The model is color-coded according to the pIDDT values (left). Ribbon representation of the model (middle) with hydrophobic residues involved in the interaction highlighted in dark green (Atg27) and red (Atg9) (right). **d**, The Atg9^CTR^-Atg27^CTR^ interaction is predicted to be stabilized by a hydrogen bond. The AF3 model as shown in Fig. 6d-e with the hydrogen bond between S802 in Atg9 and E252 in Atg27 highlighted in cyan.

**Supplementary Table 1.**
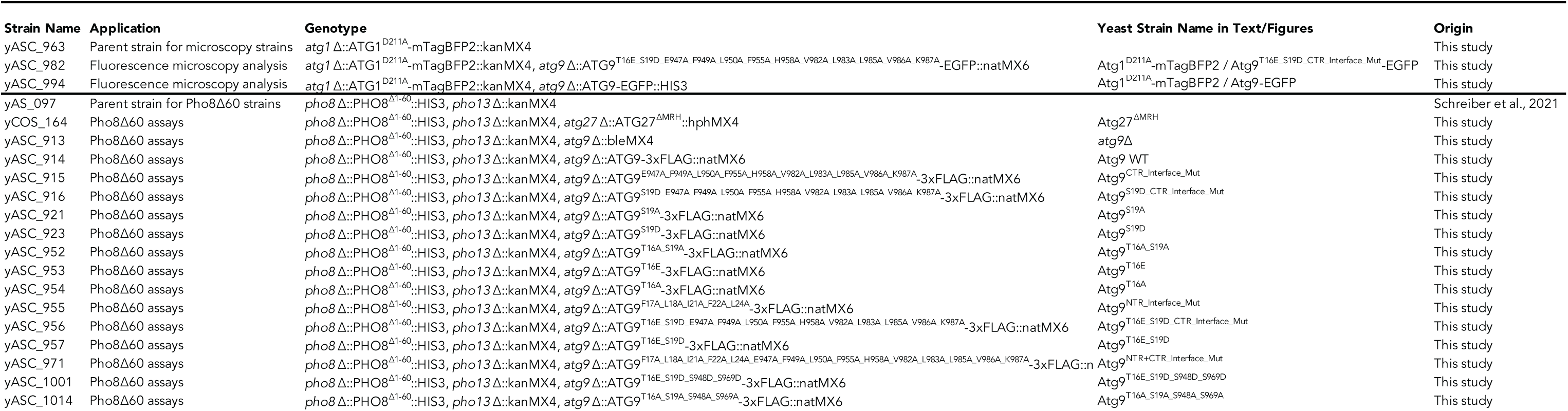
Yeast strains.

**Supplementary Table 2.**
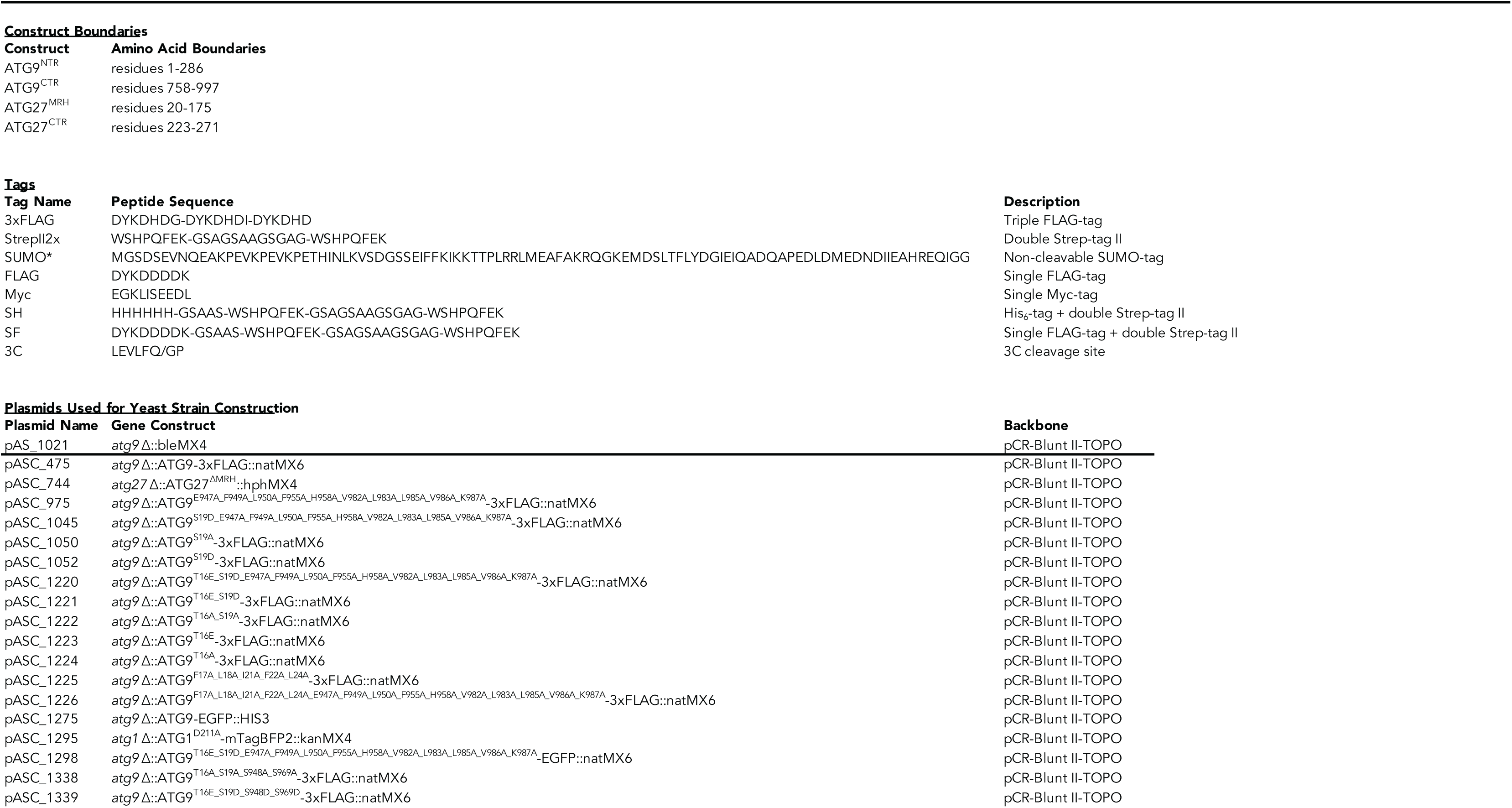
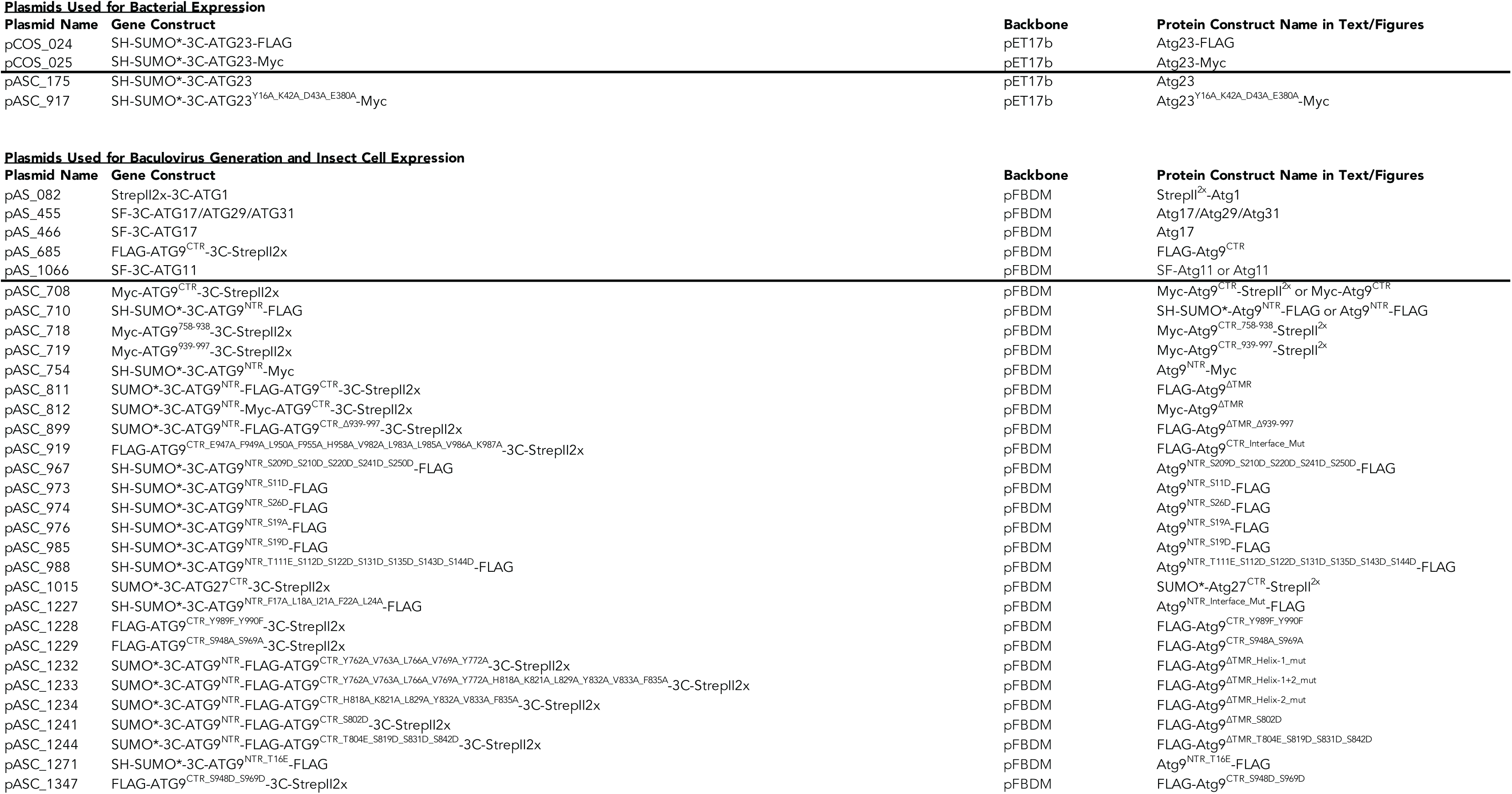
Plasmid overview.

